# The single-progenitor model as the unifying paradigm of squamous epithelial maintenance

**DOI:** 10.1101/716639

**Authors:** Gabriel Piedrafita, Vasiliki Kostiou, Agnieszka Wabik, Bartomeu Colom, David Fernandez-Antoran, Albert Herms, Kasumi Murai, Benjamin A Hall, Philip H Jones

## Abstract

Adult tissues such as the epidermis of the skin and the epithelium lining the esophagus are continuously turned over throughout life. Cells are shed from the tissue surface and replaced by cell division. Yet, the cellular mechanisms that underpin these tissues homeostasis remain poorly established, having important implications for wound healing and carcinogenesis. Lineage tracing, in which of a cohort of proliferating cells and their descendants are genetically labelled in transgenic mice, has been used to study the fate behavior of the proliferating cells that maintain these tissues. However, based on this technique, distinct mutually irreconcilable models, differing in the implored number and hierarchy of proliferating cell types, have been proposed to explain homeostasis. To elucidate which of these conflicting scenarios should prevail, here we performed cell proliferation assays across multiple body sites in transgenic H2BGFP mouse epidermis and esophagus. Cell-cycle properties were then extracted from the H2BGFP dilution kinetics and adopted in a common analytic approach for a refined analysis of a new lineage-tracing experiment and eight published clonal data sets from esophagus and different skin territories. Our results show H2BGFP dilution profiles remained unimodal over time, indicating the absence of slow-cycling stem cells across all tissues analyzed. We find that despite using diverse genetic labelling approaches, all lineage-tracing data sets are consistent with tissues maintenance by a single population of proliferating cells. The outcome of a given division is unpredictable but, on average the likelihood of producing proliferating and differentiating cells is balanced, ensuring tissue homeostasis. The fate outcomes of sister cells are anticorrelated. We conclude a single cell population maintains squamous epithelial homeostasis.

## Introduction

The squamous epithelia that cover the external surface of the body and line the mouth and esophagus consist of layers of keratinocytes. In the mouse epidermis and esophagus cell division is confined to the deepest, basal cell layer (FIGURE 1A). On commitment to terminal differentiation, proliferating cells exit the cell cycle and migrate to the suprabasal cell layers, undergoing a series of biochemical and morphological changes before being ultimately shed from the tissue surface. Cellular homeostasis requires that cells are generated by proliferation at the same rate at which they are shed. Further, to maintain a constant number of proliferating cells, on average each cell division must generate one daughter that will go on to divide and one that will differentiate after first exiting the cell cycle. However, the nature of the dividing cell population has been subject to controversy, revolving around whether dividing cells are a single, functionally equivalent population or a hierarchy of two or more distinct cell types with different proliferative potential ^1–5^. Resolving proliferating cell behavior is key for understanding not only normal tissue maintenance but also processes such as wound healing and the accumulation of somatic mutations in normal tissues during aging and carcinogenesis ^6, 7^.

**Fig. 1.**
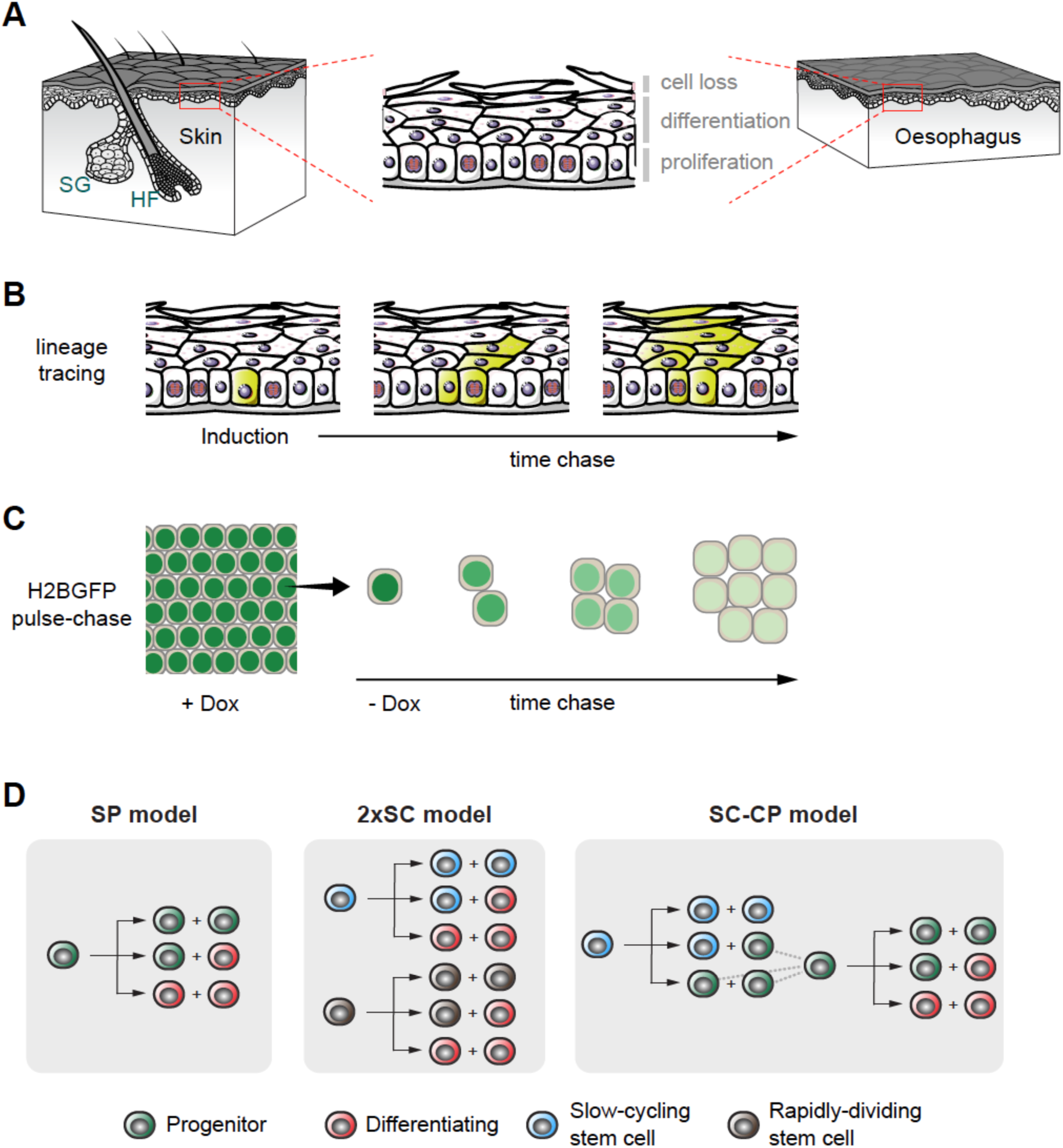
Quantitative approaches to cell behavior in murine epithelium. (A) Structure of the stratified squamous epithelia from the interfollicular epidermis (skin) and esophagus of adult mice. Proliferating keratinocytes are located in the basal layer. Upon differentiation they migrate through suprabasal layers until they are ultimately lost by shedding. A balance should be established between cell division and cell loss to guarantee tissue homeostasis. HF, hair follicle; SG, sebaceous gland. (B) Rationale of genetic lineage tracing. Low-dose induction in transgenic mice allows recombination and conditional labelling of punctuated keratinocyte progenitors in the basal layer. These cells and their progeny remain labelled and can be tracked to study clonal dynamics over time. (C) Rationale of Histone 2B-GFP (H2BGFP) dilution experiments (a top-down view of the basal layer plane is sketched). Transgenic-mouse keratinocytes express H2BGFP protein while on doxycycline (Dox) treatment. After Dox withdrawal cycling cells dilute their H2BGFP content with every division, allowing to study cell proliferation rate. (D) Different stochastic cell proliferation models invoked to explain epithelial self-renewal. Branches reflect different possible fates for a given proliferating cell upon division. SP: single-progenitor model. 2xSC: two stem-cell model, involving two independent types of proliferating cells dividing at different rates. SC-CP: stem cell-committed progenitor model, involving slow-cycling stem-cells underpinning a second population of quickly-dividing progenitors.

Whilst murine epidermis and esophageal epithelium share the same basic organization, there are significant differences between the tissues. The esophageal epithelium is uniform, with no appendages, while the epidermis is punctuated by hair follicles and sweat ducts, which form distinct proliferative compartments independent of the epidermis (FIGURE 1A) ^5, 8–10^. The structure of the epidermis also varies with body site. In typical mouse epidermis, such as that on the back (dorsum), hair follicles are frequent but there are no sweat glands ^10^. In contrast, in the mouse paw epidermis hair is absent but sweat ducts are common ^10, 11^. The ear epidermis is different again; it has uniform columns of differentiating cells, not present elsewhere ^12^. Finally, the mouse tail has the most unusual structure, being a scale forming epidermis like that of chicken legs and *Crocodillia* rather than typical mammalian skin ^13–15^. This structural diversity has motivated a range of studies to define the properties of proliferating cells at each site.

Genetic lineage tracing in transgenic mice has emerged as a powerful technique for tracking the behavior of cells within tissues (FIGURE 1B) ^16^. This is performed in mice expressing two transgenic constructs (FIGURE S1A). The first is a genetic switch, using a bacterial recombinase enzyme *Cre*, expressed either from a transgenic promoter or targeted to a specific gene^17^. A variety of *Cre* expressing mouse strains have been used for studies of esophageal epithelium and epidermis (FIGURE S1A). *Cre* is fused to a mutant hormone receptor so it is only active following treatment with a drug, giving control over when recombination is induced. Using low doses of inducing drug allows the labelling of scattered single cells. The second construct is a reporter, such as a fluorescent protein, typically targeted to the ubiquitously expressed *Gt(ROSA)26Sor* (*Rosa26*) locus. The reporter is only expressed following the excision of a “stop” cassette by *Cre*. As *Cre* mediates a genetic rearrangement, the progeny of the labelled cell also express the reporter. If cells are labelled at a low enough frequency, single-cell-derived clones of reporter expressing cells result. If a representative sample of proliferating cells is labelled and their progeny tracked over a time course, statistical analysis of the evolving clone size distributions may be used to infer cell behavior ^3^.

Alongside lineage tracing, a complementary transgenic assay may be used to detect cells cycling at different rates and infer the average rate of cell division (FIGURE 1C). This uses a transgenic, drug regulated synthetic promoter to control expression of a protein comprising Histone2B fused to green fluorescent protein (H2B-GFP) (FIGURE S1B). The H2B-GFP is initially expressed at high levels in keratinocytes. Its transcription is then shut off and levels of H2B-GFP protein measured by microscopy or flow cytometry. The stable H2B-GFP protein is diluted by cell division, so if the tissue contains cell populations dividing at different rates, the more slowly dividing cells will retain higher levels of protein ^18^. Measurements of the rate of loss of fluorescence have been used to estimate the rate of cell division ^4, 5, 19^.

Lineage tracing has ruled out older deterministic models of a proliferative hierarchy of asymmetrically dividing stem cells generating ‘transit amplifying’ cells that undergo a fixed number of divisions prior to differentiation ^3^. These models predict that clone sizes will rise and then remain stable. In multiple lineage tracing experiments, however, mean clone size has been found to increase progressively with time. However, several mutually incompatible models in which proliferating cells have stochastic fate have been proposed that do appear consistent with the data in one or more experiments (FIGURE 1D; **Suppl. Methods**).

The simplest stochastic model, the single progenitor (SP) hypothesis, proposes that all dividing keratinocytes are functionally equivalent and generate dividing and differentiating daughters with equal probability ^3, 5^. An alternative stem cell-committed progenitor (SC-CP) paradigm, applied to the epidermis proposes a hierarchy of rare, slowly cycling stem cells which generate stem and progenitor daughters. The progenitors are biased towards differentiation so continual stem cell proliferation is required ^4, 20^. A third model argues that two independent populations of stem cells (2xSC) dividing at different rates exist in the epidermis ^19^. These models all give comparably good fits to the results from individual experiments. However, each has been proposed on the basis of distinct data sets analyzed by different inference and fitting procedures, with limited testing of alternative hypotheses.

Motivated by the disparity of the proposed models of cell dynamics we set out to determine whether a consistent quantitative approach to lineage tracing data, further constrained by cell proliferation analysis in transgenic H2B-GFP mice could identify a single model that is consistent across multiple data sets in the esophagus and different epidermal regions. Here we develop an integrative stochastic-modelling approach where cell-cycle properties are extracted from new H2B-GFP dilution data, embodied in model simulations and lineage tracing results fitted, by maximum likelihood parameter inference. We test this method on a new experimental data set from mouse esophagus, and then apply it to published data sets in different skin territories, performing new H2B-GFP dilution measurements to constrain model fitting at each body site. We find that the data sets are consistent with a common, simple SP model of homeostasis. We are able to improve the parameter estimates within the model to show that the fates of pairs of sister cells are anti-correlated, and that the basal layer contains a substantial proportion of cells which will differentiate rather than going on to divide.

## Results

### Cell division rates and cycle times in epidermis and esophagus

Analysis of cell proliferation in epithelia offers a simple way to test the predictions of the disparate models of epithelial homeostasis by revealing the level of heterogeneity in the division rate of basal-layer cells. The SP model predicts a single cell population dividing at the same average rate while the alternative hypotheses argue for discrete populations dividing at different average rates. We therefore investigated the dilution of H2B-GFP in the epidermis and esophagus of *R26^M2rtTA^/TetO-H2BGFP* mice (Fig. 2A). The animals were treated with doxycycline (Dox) for 4 weeks to induce H2BGFP expression. Dox was then withdrawn and H2BGFP protein levels in individual basal keratinocytes tracked by direct measurement of GFP fluorescence using confocal imaging in epithelial wholemounts at multiple time points. We examined esophagus and epidermis from paw, ear, and tail (Fig. 2B; Fig. S2; Fig. S3). Non-epithelial cells in the form of CD45^+^ leukocytes, which retain high levels of H2BGFP, were excluded from the analysis, but served as internal reference for label retention ^5^ (e.g. Fig. 2B, **insert; Table S1**). In addition, for the analysis below we included a recently published data set from dorsal epidermis performed using an identical protocol ^21^ (Fig. S2).

**Fig. 2.**
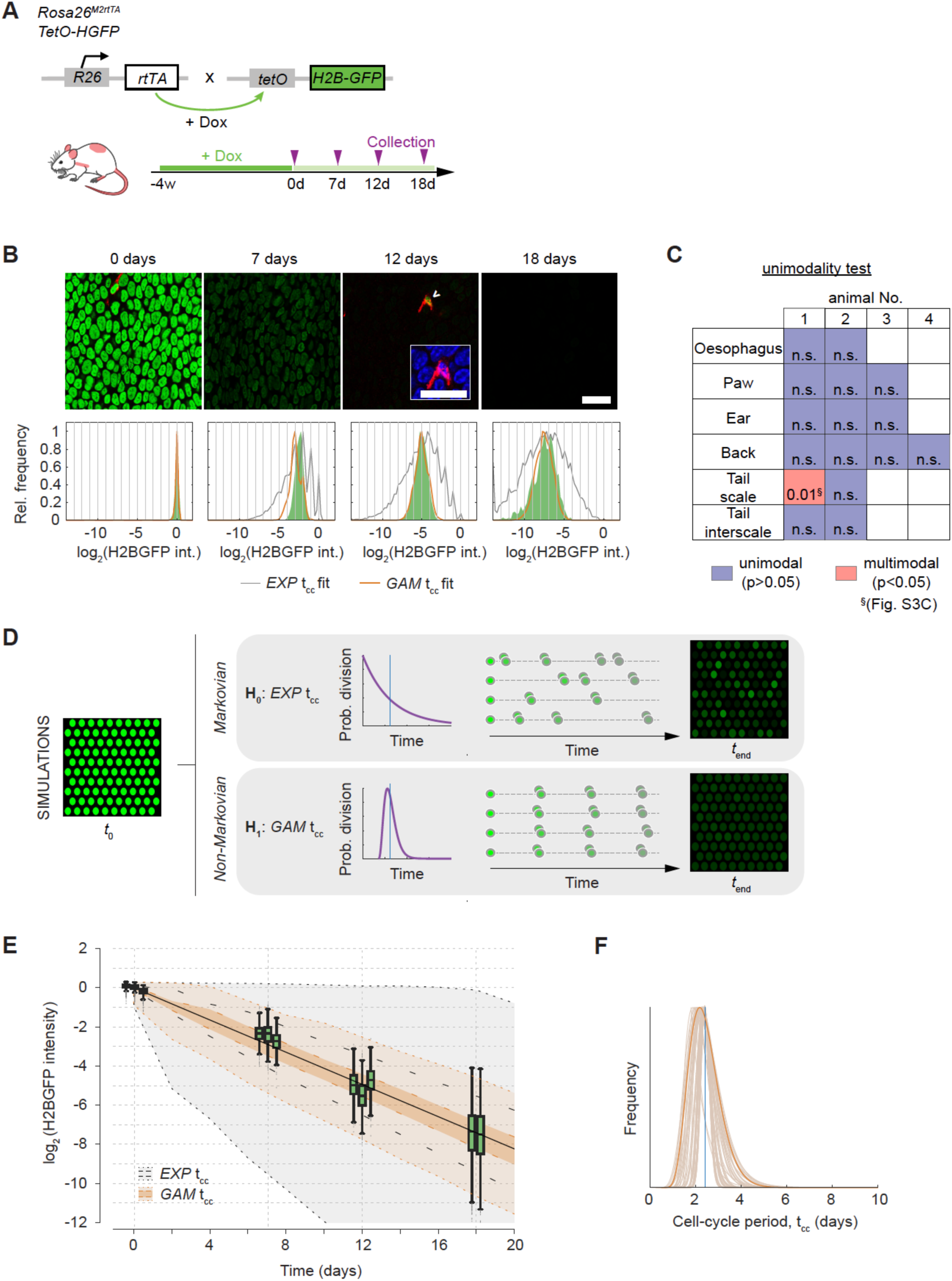
Analysis of cell proliferation in epidermis and esophageal epithelium. (A) Protocol used in H2BGFP dilution experiments: *R26^M2rtTA^/TetO-H2BGFP* mice were treated with doxycycline (Dox) to induce H2BGFP expression (green). Following Dox withdrawal, H2BGFP transcription ceases and protein levels dilute with cell division. (B) Representative confocal z stacks of the esophageal basal layer showing H2BGFP (green) and immunostaining for pan-leukocyte marker CD45 (red). Infrequent label-retaining cells (LRCs) (arrowhead) are positive for CD45 (insert; blue: DAPI). Scale bars, 20 μm. Quantifications of individual-keratinocyte H2BGFP intensity levels are presented as histograms for each time point (in green; bottom panels). Best fits for the SP model with exponential- (grey) or gamma-distributed cell-cycle periods (orange lines) are shown. (C) Outcome of Silverman’s unimodality test applied to individual-cell H2BGFP distributions at 18 days of chase in the esophagus and the different skin regions (analyses are separated per animal; no multiple-testing corrections are made on *p* values) ^35^. A single tissue in a single animal was found to be bimodal, in this case due to variability between fields of view where cells differed by a single round of division (see Fig. S3C). (D) SP-model simulations used for fitting H2BGFP intensity distributions. Taking the experimental H2BGFP intensity distributions at time 0 as input, we simulated the kinetics of cell proliferation along with H2BGFP dilution. Different forms for the distribution of individual cell-cycle times t_cc_ were considered (*EXP*: exponential; *GAM*: Gamma) for the same average division rate (blue vertical line), the narrower Gamma-shaped distributions predicting a more homogeneous dilution across keratinocytes. (E) Time course in the H2BGFP intensity distributions from esophageal epithelium (normalized to average keratinocyte intensity at time 0). Green boxplots: experimental data per mouse (computed average division rate <λ> = 2.9/week; solid black line). Grey region: range of H2BGFP intensities predicted from models assuming cell cycle periods were highly heterogeneous, i.e. exponentially distributed (interquartile range given by inner dashed black lines). Light orange region: range of H2GFP intensities inferred with a gamma cell cycle period (interquartile range in dark orange, delimited by inner dashed orange lines). (F) Most-likely (gamma) shapes for the distribution of the cell-cycle period of esophageal keratinocytes, as estimated from fits to the H2BGFP dilution data. A conservative solution (in dark orange) is used for further inference. Vertical blue line: average cell cycle period.

We first examined images for the presence of label retaining cells (LRCs) (**Table S1**). We found no keratinocyte LRC in the basal cell layer of the esophagus or any epidermal site other than the interscale region of the tail (Fig. S3A). Rare keratinocyte LRCs (4/1923, i.e. 0.2% of basal layer keratinocytes) were observed in interscale epidermis, in a single animal, 18 days after DOX withdrawal. Their scarcity however suggests that they are unlikely to make a substantial contribution to tissue maintenance.

Next, we performed a quantitative analysis of the time series of the individual-cell H2BGFP intensity histograms (**Table S2**). If there were multiple subpopulations of cells proliferating at different rates the distribution of H2BGFP intensities would progressively diverge, becoming wider over time. We found no evidence of such behavior in the esophagus and at multiple sites in the epidermis (Fig. 2B; Fig. S2). Specifically, several statistical tests of the modality of the distribution were applied, showing no evidence for multiple populations (Fig. 2C; Fig. S3; **Table S3; Suppl. Methods**).

To further challenge the hypothesis that there is a single proliferating cell population, we examined whether this model can recapitulate the observed H2BGFP intensity distributions at each time point. For a given *average* division rate, we performed simulations of H2BGFP-dilution kinetics under a wide range of possible underlying (Gamma) distributions for individual cell-cycle times (Fig. 2D; **Suppl. Methods**). We find that the form of the H2BGFP histograms over time can indeed be fully described by a single population of cells, dividing within a relatively narrow range of cell cycle times, further supporting the SP model (Fig. 2B,E,F; Fig. S2; **Table S4**).

Altogether, these observations strongly argue against scenarios of heterogeneous proliferating cell populations, such as the SC-CP or 2xSC models, at all sites other than in the tail where marked variation between animals precluded reliable inference on cell proliferation rates (Fig. S3). We conclude that in the basal layer of the epidermis at multiple body sites and in the esophagus proliferating cells divide at a unique average rate with highly homogenous cell-cycle periods, consistent with the SP model (**Table 1**).

### A common analytical approach integrating cell proliferation and lineage tracing data

The ability of lineage tracing to track the behavior of cohorts of proliferating cells and their progeny over time courses extending to many rounds of cell division offers the potential to validate models of homeostasis. Having established the homogeneity in the division rate of basal-layer cells, we then set out to determine whether clonal dynamics across different lineage tracing data sets were consistent with the SP paradigm.

Multiple lineage tracing studies have been published but these used distinct approaches to infer models of cell behavior and did not apply the additional constraint imposed by measuring the cell cycle time distribution ^3, 4, 11^. Computational simulations revealed that the SP, SC-CP and 2xSC models all predict very similar development of clonal features over time, which rendered them hardly distinguishable from lineage tracing data alone (**Suppl. Methods**). However, as our cell-proliferation analyses do not support the SC-CP and 2xSC paradigms we focused on testing the SP model. By incorporating the measurement of the *average* division rate, we could reduce the uncertainty in the parameter estimation, a problem that has been generally overlooked in these stochastic models (Fig. S4). In turn, whilst long-term model predictions on clone-size distributions remained largely unaffected by the assumptions on the cell-cycle time *distribution*, introducing realistic estimates for the distribution of individual cell-cycle lengths affected short-term clone-size predictions, impacting on the inferred parameter values (Fig. S5). This disputes most common, Markovian implementations that take the biologically implausible assumption that cell cycle times are distributed exponentially (i.e. the likeliest time for a cell to divide is immediately after the division that generated it). We therefore developed a robust quantitative approach where cell-cycle attributes estimated from H2B-GFP experiments were embodied in (non-Markovian) model simulations, and a subsequent maximum likelihood inference (MLE) method was applied across the available data sets for each body site to challenge whether each of them was consistent with the SP paradigm (**Suppl. Methods**).

### Clonal dynamics in esophageal epithelium

In order to explore *in vivo* clonal dynamics, we began by studying a new lineage tracing dataset in mouse esophageal epithelium (Fig. 3A). In this experiment we used a strain (*Lrig1-cre)* in which a tamoxifen-regulated form of *Cre* recombinase and enhanced green fluorescent protein (EGFP) are targeted to one allele of the *Lrig1 locus* ^8, 22–24^. We found LRIG1 protein was ubiquitously expressed in the basal layer of esophageal epithelium in wild type mice (Fig. S6A). Consistent with this finding, in *Lrig1-cre* animals, EGFP, reporting *Lrig1* transcription was detected in 94+/-0.3% (s.e.m.) of basal cells (Fig. S6B). These observations indicate *Lrig1* is widely expressed in the proliferative compartment of esophageal epithelium and is suitable for lineage tracing of proliferating esophageal keratinocytes (**Suppl. Methods**).

**Fig. 3.**
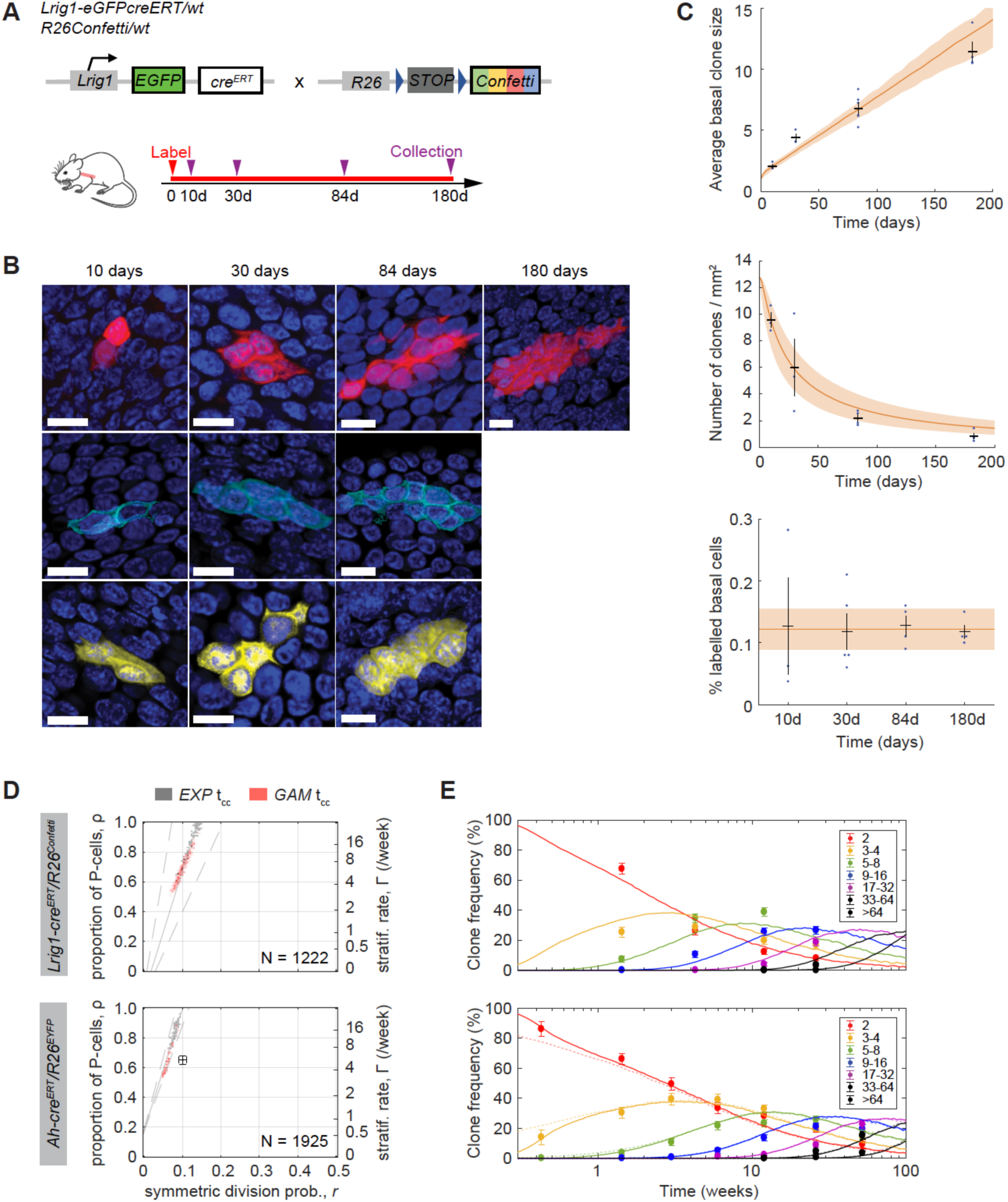
Quantitative lineage tracing in esophageal epithelium. (A) Protocol: clonal labelling was induced in *Lrig1-eGFPcre^ERT/wt^ R26^flConfetti/wt^* mice and samples analyzed at different times from 10 days to 180 days post-induction, as single labelled cells develop into clones. (B) Rendered confocal z stacks of the esophageal basal layer showing typical clones at the times indicated. Red is RFP, light blue is CFP, yellow is YFP, blue is DAPI. Scale bars, 10 μm. (C) Quantitative traits of the labelled clone population over time: average basal-layer clone size (i.e. mean number of basal cells/surviving clone) (top panel), average density of labelled clones in the basal layer (middle panel), average fraction of labelled basal cells at the indicated time points (bottom panel). Observed values (blue dots) and error bars (black; mean ± s.e.m.) from n ≥ 3 animals. Orange lines: SP model fit (shaded area corresponds with 95% plausible intervals). Orange line and shading in last panel show mean and s.e.m. across all time points, which is consistent with homeostatic behavior. (D) SP model parameter inference on *Lrig1*- and *Ah*-*Cre^ERT^* driven lineage tracing datasets from esophagus (left panels) ^5^. Parameter estimates are affected by the underlying modelling assumptions on the cell-cycle period, whether default exponential cell-cycle time distributions were considered (solutions in grey) or realistic gamma distributions implemented, as inferred from the cell proliferation analysis (solutions in orange). Regions within the dashed grey lines fall consistent with the predicted ρ/*r* ratios from the linear scaling of the average clone size. The total number of clones counted in each data set is displayed in the corresponding graph and previous parameter estimates given in ^5^ shown as black error bars. Experimental *Lrig1*- and *Ah*-*Cre^ERT^* derived basal-layer clone sizes (right panels; dots with error bars as standard error of proportion) are excellently fit with the SP model with gamma-distributed cell cycle times (lines; prediction from MLE). Dim dashed lines: fits from ^5^. Frequencies for each clone size (basal cell number) are shown in different colors.

To track the fate of basal cells, *Lrig1-cre* mice were crossed with the *Rosa26^flConfetti/wt^* (*Confetti*) reporter strain which labels cells with one of four possible fluorescent proteins (green, GFP, cyan, CFP, yellow, YFP or red RFP) after recombination (Fig. S6C-D) ^25, 26^. In some *Cre* inducible mouse lines, reporter expressing clones can appear without induction with Tamoxifen. However, no fluorescent protein expression was found in adult uninduced *Lrig1-cre/Confetti* mice (**Table S5**) ^27^. Next, cohorts of *Lrig1-cre*/*Confetti* animals were treated with a low dose of Tamoxifen that resulted in labelling of only 1 in 300 ± 106 (mean ± SEM) basal cells at 10 days post-induction. Clones containing one or more basal cells were imaged in esophageal epithelial wholemounts from at least 3 mice at multiple time points over 6 months following induction (Fig. 3B; **Table S5**). Only CFP, YFP and RFP expressing clones were counted because of *Lrig1* driven GFP expression in all basal cells.

The clone data set displayed several important features. The density (clones/area) of labelled clones decreased progressively, consistent with clone loss through differentiation, while the number of basal and suprabasal cells in the remaining clones rose (Fig. 3C; Fig. S7A). The proportion of labelled basal cells remained constant during the experiment, indicating the labelled population was self-maintaining over a six month period, consistent with labelled cells being a representative sample of all proliferating cells in the homeostatic tissue (Fig. 3C). At late time points, the clone size distribution scaled with time (i.e. the probability of seeing clones larger than x times the average clone size became time-invariant, following a simple exponential f(x) = e^-x^) (Fig. S7B). Collectively, these features are hallmarks of neutral competition, in which clonal dynamics result from stochastic cell fates, with an average cell division generating one proliferating and one differentiating daughter cell, a scenario consistent with the SP model (**Suppl. Methods**) ^3, 28^.

The measurement of the average cell division rate (λ) and inference of the cell cycle time distribution constrain the fitting of lineage tracing data, providing a stringent test of the candidate SP model (Fig. S4B-C; Fig. S5E). Within this paradigm unknown parameters are the probability of a progenitor cell division generating two dividing (PP), or two differentiating (DD) daughters (*r*), and the stratification rate (Γ), which in homeostasis sets the fraction of progenitor cells in the basal layer (ρ) (Fig. S4A). Our technique for identifying the most appropriate cell-cycle distribution coupled with a MLE grid search discriminated parameter estimates that gave excellent fits on clone-size distributions at both early and late time points for the *Lrig1/confetti* dataset (Fig. 3D-E; **Table 1; Table S4**).

Next, we applied the same approach to an independent, published lineage tracing data set from esophageal epithelium where clones were labelled with YFP by *Cre* expressed from an inducible *Cyp1a1* (*Ah*) promoter in *AhYFP* mice ^5^. A close fit to the SP model was obtained for very similar parameter values to the ones above, with cell-cycle time constrains resulting in an improved match over short-term clone sizes compared with fits in the original publication (Fig. 3D-E; Fig. S7C; **Table 1; Table S4**). As a further validation, we tested our parameterized model against a third, more limited dataset from *Krt15-cre^PR1^ R26^mT/mG^* mice in which a red-to-green fluorescent reporter was used with inducible *Cre* expressed from a *Krt15* promoter (**Suppl. Methods**) ^29^. Although the SP paradigm was criticized by these authors, it yielded an adequate fit with their own data over the experimental time course (Fig. S7D; **Table S4**). The consistent fit of the SP model to three independent lineage tracing data sets using different combinations of transgenic *Cre* and reporter alleles strongly supports the conclusion that the esophageal epithelium is maintained by a single progenitor population and reinforces the reliability of our parameter estimates.

### Clonal dynamics in skin epidermis

We next investigated clonal dynamics in the epidermis through available lineage tracing data sets from the typical interfollicular epidermis of the mouse hind paw, ear and back (dorsum). Applying the MLE approach constrained by the cell-cycle time analysis at each body site yielded improved fits of the SP model to data from paw epidermis *in Axin2-cre^ERT^R26^Rainbow^* animals and from ear and dorsal interfollicular epidermis in *AhYFP* mice compared with those fits reported in the original publications (Fig. S8; **Suppl. Methods**) ^11, 21, 30^. Despite differences in average keratinocyte division rates across territories (λ ≈ 2.0, 1.5, 1.2/week for hind paw, ear and dorsum, respectively), all analyzed regions share comparable intermediate proportions of progenitor basal cells ρ (∼55%, the rest corresponding to differentiating basal cells) and a predominance of asymmetric cell divisions (i.e. low inferred values for the probability of symmetric division, *r* < 0.25) (**Table 1; Table S4**).

Particularly relevant are the implications for the mode of keratinocyte renewal in back skin, as a previous work claims that two stem cell populations dividing at different rates coexist at this site (2xSC model)^19^. This argument was supported by a quantitative analysis of H2BGFP dilution patterns in *Krt5^tTA^/pTRE-H2BGFP* mice, a system that differs from that we use above in that mice are treated with Dox to suppress H2B-GFP expression instead of using it as activator (Fig. S9A)^19^. However, in that publication, in the rejection of the SP model, exponential distributions for cell division/cell stratification rates were assumed, which we have here shown to be inappropriate for the short time scale of the experiment (**Suppl. Methods**). Our computational reanalysis, constrained by cell-cycle time distributions demonstrated the SP model gave as good a fit to the *Krt5^tTA^/pTRE-H2BGFP* dilution data as the more complex 2xSC hypothesis (Fig. S9B; **Suppl. Methods; Table S4**). Indeed, the inferred parameter values from the *AhYFP* mouse back skin epidermis proved robust, providing good fits to another lineage tracing data set from the same body site in *Lgr6-eGFPcre^ERT^Rosa26^flConfetti^* mice (Fig. S9C; **Suppl. Methods; Table S4**) ^33^.

Finally, we turned to revisit clonal dynamics in the mouse tail epidermis. Previous studies of tail have argued that the hierarchical SC-CP paradigm applies to proliferating cells in the interscale areas while the SP paradigm describes behavior in the scale regions (Fig. S3A) ^4, 20^. These claims were primarily supported by the observation of LRCs in the interscale region in H2BGFP dilution experiments in *Krt5^tTA^/pTRE-H2BGFP* mice. However, our quantitative reanalysis of this data set revealed the SP model fits the reported H2BGFP intensity histograms over time as well as the SC-CP model (Fig. S10A; **Table S4**)^20^. Even though we cannot discard the possibility of a subpopulation of slow-cycling stem cells in the tail, such cells would seem to be rare in interscale epidermis (**Table S1**). Further analysis argued that there was no conflict between the reported tail lineage tracing data and the SP model (Fig S10B-H; **Suppl. Methods; Table S4**).

## Discussion

Overall, we find that combining cell cycle distribution analysis with lineage tracing argues mouse esophageal epithelium and epidermis are maintained by a single population of progenitor cells, with the sole possible exception of the inter-scale compartment of tail skin. The quality of the fit of the SP model to the data is equivalent to or exceeds that of more complex models, rendering the need to invoke additional cell populations redundant. The nine lineage tracing data sets analyzed include a variety of *Cre* and reporter strain combinations, and are all consistent with the SP model. In addition, live imaging studies of the epidermis are consistent with a single proliferating cell population maintaining the tissue ^31^.

Quantitative analysis of cell proliferation in the different tissue types reveals further constraints that must be considered by researchers exploring the appropriateness of alternative models. The original SC-CP and 2xSC models invoked 12% and 30% of basal-layer keratinocytes constitute slow-cycling stem cells, respectively ^4, 19^. Histone dilution experiments have allowed us to make strong statements about the nature of any proposed second population. For each body site, with the exception of tail inter-scale epidermis, no keratinocyte label retaining cells were detected in over 2000 cells imaged at each location. It follows that any slow cycling stem-cell population must have substantially fewer than one slow-cycling cell per thousand basal keratinocytes to be compatible with observations reported here, making it unlikely that such slow cycling cells will make a detectable contribution to tissue homeostasis. The hypothesis that two subpopulations exist, but that they both divide at a similar rate, is hard to sustain in the face of the close agreement of the simpler SP model across all the analyzed data sets.

The improved resolution of parameter estimates reveals differences in cell division rates across the epidermis and the esophagus (**Table 1**). Proliferating cells divide rapidly in the esophageal epithelium, on average every ∼2.4 days (similar to keratinocyte turnover rate in oral mucosa), while progenitor cells cycle comparatively slower in the epidermis, on average between 3.5-6 days depending on body site ^32^. However, our study suggests individual cell-cycle periods are tightly controlled, showing little variation around average division rate, per territory.

The proportion of progenitor cells in the basal layer and the probability of symmetric cell division outcomes (r) are similar across body sites (**Table 1**). The insight that a substantial proportion of cells in the basal layer will proceed to differentiate rather than divide will be important for the interpretation for the growing body of single cell RNA sequencing data in these tissues ^32, 33^. In addition, the low values of r we identify give insight into the basis of cell fate determination (Fig. 4). In principle, if every basal cell divides or differentiates with equal probability, as proposed by Leblond, r will be 0.25, as expected from any pair of uncorrelated basal cells ^1^. However, this scenario is excluded by our analysis. Instead, the consistent values of r <0.25 indicate the fate of sister cells is preferentially anti-correlated. This phenomenon can be associated to local coordination of neighboring cell stratification and division events, as has been observed *in vivo* in adult ear epidermis by live imaging ^34^. Our results argue that anticorrelation of sister cell fates applies generally in the epidermis and esophagus, pointing to common mechanisms of keratinocyte cell fate regulation. We conclude that the SP model explains squamous epithelial homeostasis.

**Fig. 4.**
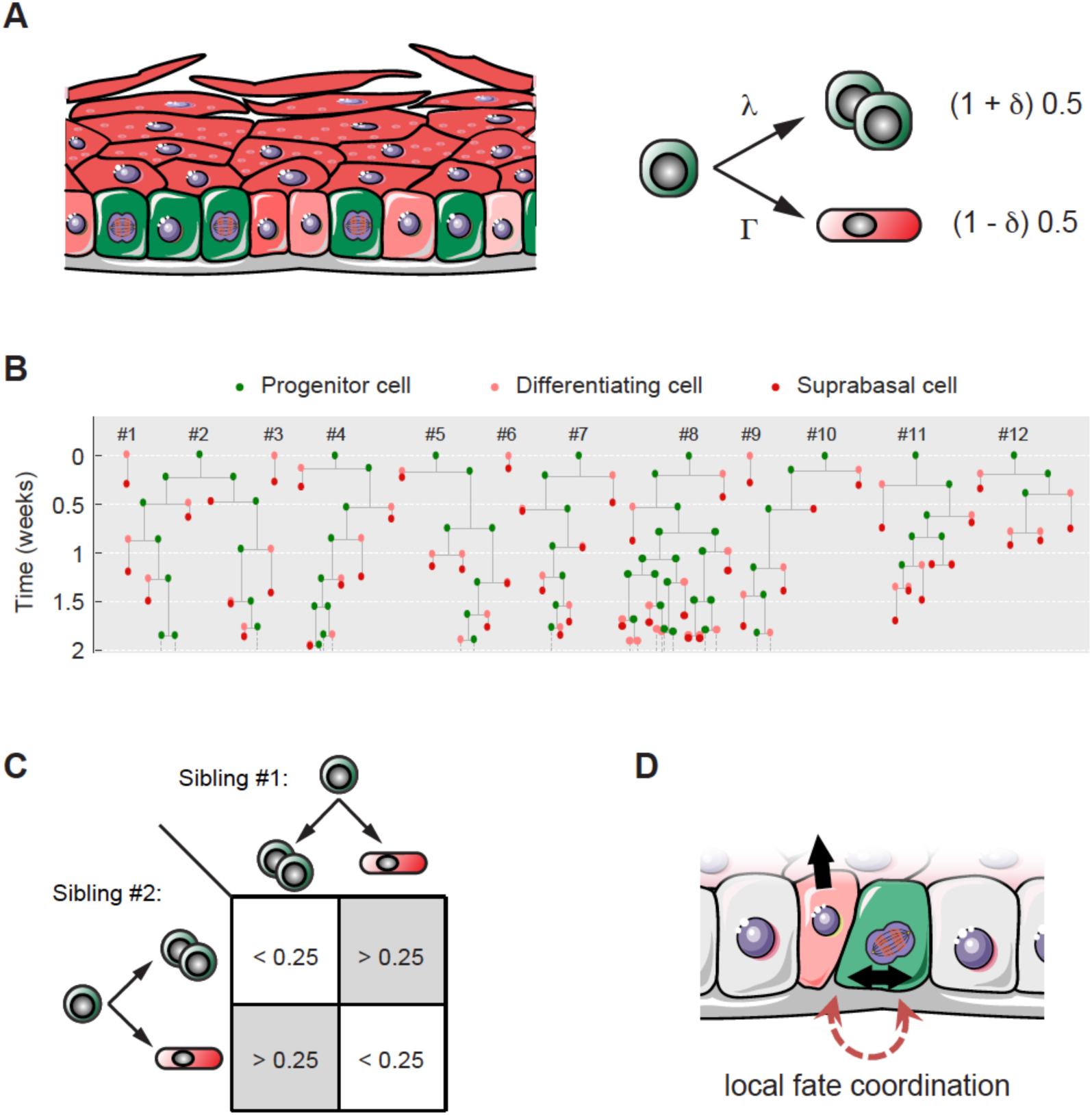
Cell fate coordination underpins single progenitor dynamics in squamous epithelia. (A) Squamous epithelial tissues are maintained by a single population of progenitor cells. Left panel: Progenitor cells (in green) share the basal layer with post-mitotic keratinocytes in early stages of differentiation (pale red), which are transiently retained in the basal compartment before stratification. Right panel: Simplified representation of the single progenitor model focusing on individual basal cell fates. Epithelial cell dynamics are dominated by stochastic but skewed fates through spatial coordination between neighboring or sibling cells. Individual basal cells undertake dichotomic decisions: they have the potential to divide but can alternatively differentiate exiting the basal layer. Both probabilities are balanced (50%-50%) across the entire homeostatic tissue, but can be decompensated or skewed for individual cells depending on their local niche, as reflected with the parameter δ (which works as a context-dependent modulatory factor). (B) Stochastic progenitor fates explain a scenario of neutral clonal competition dynamics where clones develop into heterogeneous sizes, constrained by cell-cycle time control and fate coordination effects. Displayed is a representative set of epithelial clone dynamics simulated using the parameters inferred for murine esophageal epithelium homeostasis. (C, D) Our results demonstrate that the outcome of sibling keratinocyte cells is commonly biased towards an excess of asymmetric fates where one decides to divide while the other differentiates, in agreement with a single progenitor model with low values of *r* (*r*<0.25) (see Fig. S4A, secondary processes in dim color).

## Methods

### Animals

Doubly transgenic, *Lrig1-eGFPcre^ERT/wt^ R26^flConfetti/wt^* mice were generated for lineage tracing studies in esophageal epithelium, by crossing *Lrig1-eGFP-ires-cre^ERT2^* mice ^8^ onto a *Rosa26^flConfetti^* multicolor reporter line ^25^. Transcription of the *Cre* recombinase-mutant estrogen receptor fusion protein (Cre^ERT^) is under the control of an endogenous allele of *Lrig1*. Following induction with tamoxifen, Cre^ERT^ protein internalizes into the nuclei and excises a *LoxP*-flanked “STOP” cassette resulting in the expression of one of the four Confetti fluorescent reporters (YFP, RFP, CFP or GFP). *R26^M2rtTA^/TetO-H2BGFP* mice, doubly transgenic for a reverse tetracycline-controlled transactivator (rtTA-M2) targeted to the *Rosa26* locus and a HIST1H2BJ/EGFP fusion protein (H2BGFP) expressed from a tetracycline promoter element, were used for label retaining experiments ^5, 21^. H2BGFP expression is induced by treatment with doxycycline (Dox) and dilution of H2BGFP protein content can be chased upon Dox withdrawal. All animals were induced at 8-12 weeks age. Cohorts of at least two or three animals per time point were culled and esophagus and/or skin epidermis collected for analysis.

### Wholemount preparation & immunostaining

Esophageal epithelium wholemounts for lineage tracing were prepared as follows: The esophagus was cut longitudinally and the middle two thirds of the tract was incubated for 3 hours in 5 mM EDTA in PBS at 37°C. The epithelium was then peeled away from the underlying submucosa, stretched and fixed for 30 minutes in 4% paraformaldehyde in PBS. Samples were stored in PBS at 4°C until subsequent analysis. Skin pieces of approximately 0.5cm^2^ were cut and incubated for 1 hour in 5 mM EDTA in PBS at 37°C. Skin epidermis was then peeled away using fine forceps and processed as described above for the esophageal epithelium.

For staining, wholemount samples were incubated in Permeabilization Buffer (PB) (0.5% BSA, 0.25% Fish Skin Gelatin (FSG), 0.5% Triton X-100 /PBS) for 15 minutes at room temperature (RT), then blocked in 10% goat or donkey serum/PB (according to the secondary antibody used) for 1 hour at RT and incubated overnight with primary antibody at 4°C. Primary antibodies used were: Lrig1 antibody (R&D Systems, Cat. AF3688), ITGA6 antibody (clone GoH3, Biolegend, Cat. B204094), Alexa Fluor® 647 anti-CD45 (clone 30-F11, Biolegend, Cat. 103124), Keratin 14 antibody (clone Poly19053, Biolegend, Cat. 905301). Samples were subsequently washed 4 times for 30 minutes in 0.2% Tween-20/PBS and incubated with an appropriate secondary antibody for 3 hours at RT. Secondary antibodies used were Goat or Donkey Alexa Fluor 488/546/555/647 (Molecular Probes). A washing step with 0.2% Tween-20/PBS was repeated and samples were incubated for 30 minutes with DAPI (Sigma-Aldrich) and finally mounted in DAKO Vectashield Mounting Medium with DAPI (Vector Labs).

### Dilution of Histone 2B-GFP (H2BGFP) protein content

*R26^M2rtTA^/TetO-H2BGFP* animals were treated with doxycycline (Dox, 2 mg/ml in drinking water sweetened with 10% blackcurrant & apple) for 4 weeks. Dox was then withdrawn and animals culled at different time points to track H2BGFP florescence dilution. Epithelial wholemounts from esophageal epithelium and skin epidermis were imaged on a Leica TCS SP8 confocal microscope using 20x or 40x objectives at 1024 x 1024 resolution, line average 4 and 400Hz scan speed. Individual-cell H2BGFP intensities were determined by image segmentation/nuclear identification, using a semi-automated object-recognition macro (based on the DAPI channel) built in ImageJ, and the process completed by manual curation. Per-cell intensity values given are averaged over all nuclear pixels.

### Lineage tracing

Low-frequency expression of the Confetti reporters in the *Lrig1-eGFP-ires-cre^ERT2^ R26^flConfetti/wt^* mouse esophagus was achieved by inducing 10-weeks old animals with intraperitoneal injection of a single dose of 1mg tamoxifen (100 μl of 10 mg/ml) on two consecutive days ^8^. This resulted in a labeling efficiency of 1 in 301 ± 106 (mean ± SEM) basal cells by 10 days post induction (allowing individual clone tracking without merging). Between three and six mice were culled per time point. Confocal images of immunostained wholemounts were acquired on a Leica TCS SP8 confocal microscope (x10, x20 and x40 objectives; typical settings for z-stacks acquisition: optimal pinhole, line average 4, bi-directional scan with 400-600 Hz speed, resolution of 1024×1024 pixels).

The number of nucleated basal and suprabasal cells per labelled clone was counted under live acquisition mode. GFP-labelled clones were not scored due to the difficulty of distinguishing them from the constitutive basal GFP expression driven by the Lrig1 cassette. CFP, RFP and YFP clones were pooled together for further analysis (histograms (distributions) of basal-layer clone sizes and average number of basal cells). A total of 300, 315, 302 and 305 labelled clones from 3, 3, 6 and 4 mice at 10, 30, 84 and 180 days post-induction, respectively, were quantified. Regarding the time courses in the number of clones per unit area and the proportion of labelled basal cells, only RFP clones were considered given the low, variable induction of the other florescent reporters and their overall small contribution (including these numbers did not alter the conclusions).

Lineage-tracing data from *Ah-cre^ERT^ R26^flEYFP^* derived clones in esophagus ^5^, ear ^30^ and dorsal epidermis ^21^ were obtained from experimental colleagues (data available upon request). Data on induced *Axin2-cre^ERT^ R26^Rainbow^* clones in hindpaw ^11^ and *Lgr6-eGFPcre^ERT^ R26^flConfetti^* in back epidermis ^33^ were kindly provided by the authors. Data from lineage tracing in scale and interscale tail epidermis ^20^ was accessed through the online publication material, while authors were unable to provide original data from ^4^. Data on *Krt15-cre^PR1^ R26^mT/mG^* mouse esophagus ^29^ were retrieved by digitalizing Fig. 2E and Fig. S3B from the original publication. A similar procedure was used to extract *Krt5^tTA^/pTRE-H2BGFP* dilution data from back skin (Fig. 3 from ^19^) and tail epidermis (Fig. 3K from ^4^).

### Mathematical modelling and statistical inference

For details of theoretical modelling and computational methods used to infer cell behavior and clonal dynamics, see Supplementary Methods: Theory.

Code used in computational modeling is available in Github: https://github.com/gp10/Piedrafita_etal_SI_code/

## Acknowledgements

We are grateful to Xinhong Lim and Roeland Nusse; Kyogo Kawaguchi, Allon M. Klein and Valentina Greco; Anja Füllgrabe and Maria Kasper; and David Shalloway for kindly sharing raw lineage-tracing data and computational algorithms from previous publications, and for their valuable comments. We thank María P. Alcolea for useful feedback on H2BGFP chase experiments and Kim Jensen for sharing the *Lrig1-creERT* mouse strain. We thank Esther Choolun and staff at the MRC ARES and Sanger RSF facilities for excellent technical support and Hall and Jones’ group members for critical comments and useful discussion. This work was supported by funding from core grants from the Wellcome Trust to the Wellcome Sanger Institute, 098051 and 296194, a Cancer Research UK Programme Grant to P.H.J. (C609/A17257) and the Royal Society (UF130039 to B.A.H.).

## Author contributions

GP performed the computational simulations and carried out the analysis of experimental results. AW carried out the linage tracing experiments. BC and DFA performed H2BGFP dilution experiments. AH and KM contributed with image acquisition and image analysis. VK helped implementing the Non-Markovian simulation algorithm. BAH and PHJ supervised the work. GP, BAH and PHJ designed the work and wrote the manuscript. All authors contributed in the elaboration of the final version.

## Supplementary Figures and Legends

**Fig. S1.**
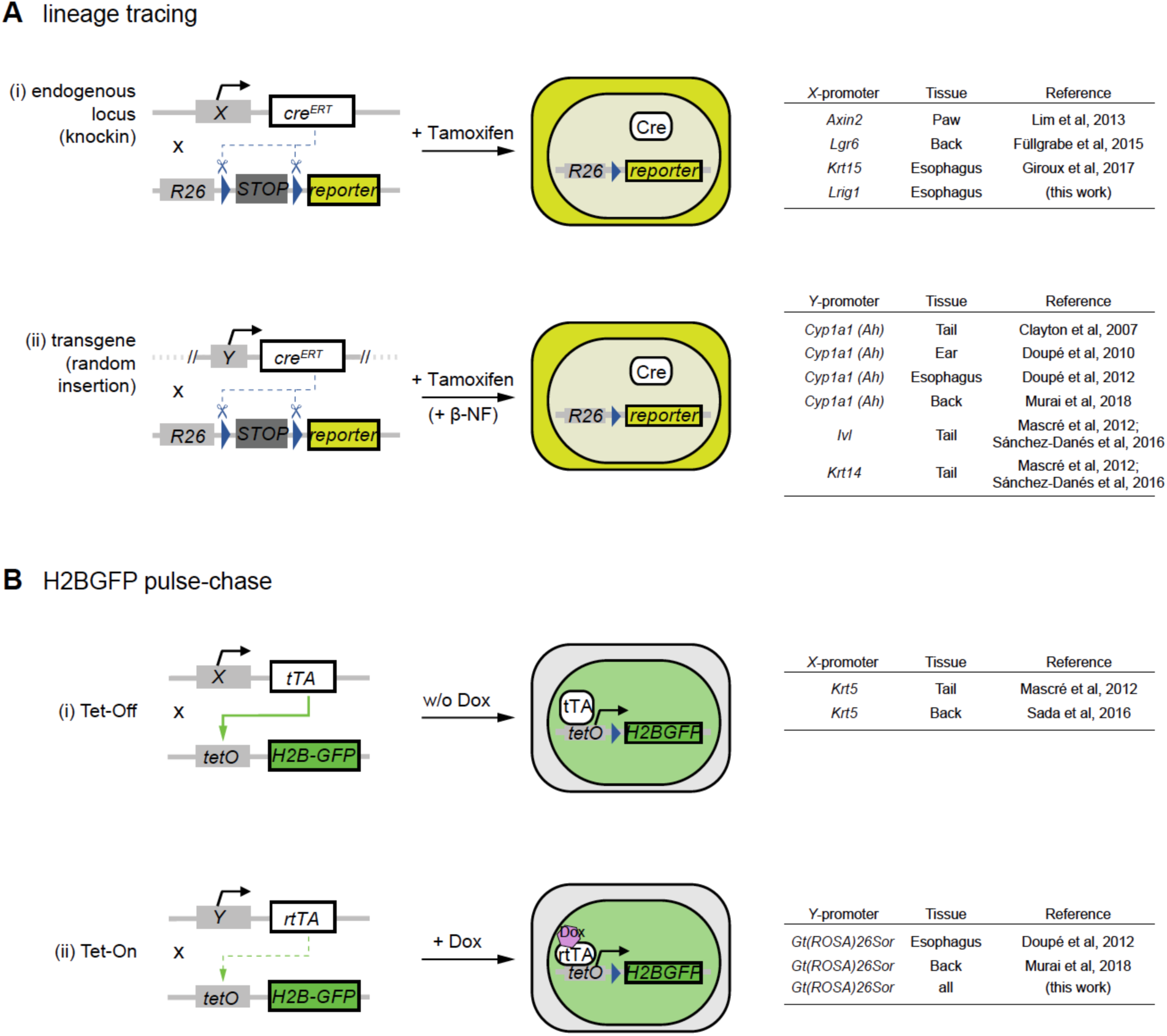
Transgenic mouse models used for lineage tracing and cell-proliferation studies. (A) Transgenic mice for lineage tracing are designed with two genetic constructs. The first codes for a bacterial *Cre* recombinase-mutant estrogen receptor fusion protein (CreERT), which can be targeted to a specific endogenous locus (i) or be under control of a transgenic promoter, randomly inserted in the genome (ii). The second construct codes for a conditional fluorescent protein reporter, typically targeted to the ubiquitously expressed *Rosa26* locus. Treatment with tamoxifen induces *C*re protein internalization to the nucleus, allowing expression of the reporter following Cre-mediated excision of a *loxP*-flanked STOP cassette. Specific details of constructs used in previous literature for lineage tracing in squamous epithelia are shown on the right (Tables). Note expression of *Cre* from the transgenic *Cyp1a1* arylhydrocarbon receptor, *Ah*) promoter requires additional treatment with a *Ah* inducer, β-napthoflavone (β-NF). (B) Transgenic mice for H2BGFP dilution experiments are designed with a first construct, typically targeted to a constitutive promoter, coding either for a tetracycline-controlled transactivator (tTA; Tet-Off system) (i) or a reverse tetracycline-controlled transactivator (rtTA; Tet-On system) (ii). A second construct codes for a Histone 2B-green fluorescent protein fusion (H2BGFP) controlled by a tetracycline-response promoter element (*tetO*; sometimes referred to as *pTRE*). Treatment with tetracycline or its derivative doxycycline (Dox) preempts tTA protein from binding to *tetO* elements in Tet-Off systems, causing repression of *pTRE*-controlled H2BGFP expression, whilst it is required for binding of rtTA to *tetO* elements in Tet-On systems, hence having an opposite effect. Dox is administered for induction and withdrawn during the H2BGFP dilution chase in Tet-On mice, while in Tet-Off animals its application gets required for the duration of the experiment.

**Fig. S2.**
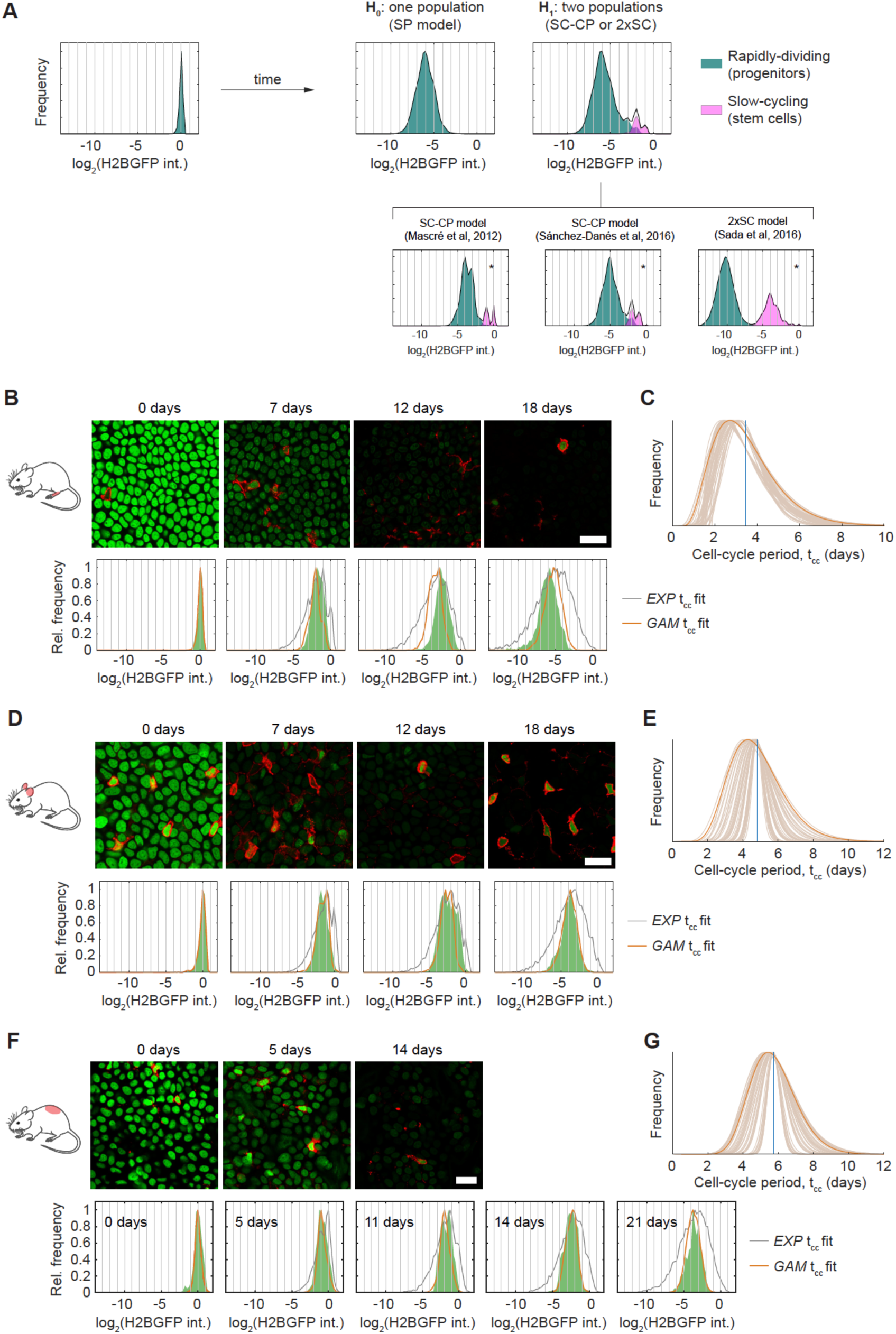
H2BGFP dilution analysis and keratinocyte cell cycle inference in different skin territories. (A) Theoretical distributions of individual-cell H2BGFP intensities expected after 3 weeks of chase under each of these hypotheses: a single population of progenitor cells cycling at a similar pace (SP model), or two populations of cells cycling at different rates (SC-CP or 2xSC scenarios), assuming realistic, gamma-distributed cell-cycle periods for either population. Notice that each indent of a jagged pattern corresponds with a different number of cell divisions (i.e. a unit in the normalized log_2_ scale). By default, simulations considered an average division rate for stem cells 4x slower than for progenitors, i.e. λ_P_ = 2/week, λ_SC_ = 0.5/week (top panels). Predictions under the specific paradigm and parameter conditions invoked in some previous work are shown below. All theoretical SC-CP or 2xSC scenarios represented in this figure were detected as significant by 6 different unimodality tests. (B, D, F) Representative confocal z stacks of hindpaw (plantar), ear and dorsal epidermal basal layer, respectively, from *R26^M2rtTA^/TetO-H2BGFP* mice, showing H2BGFP (green) and immunostaining for pan-leukocyte marker CD45 (red). Label-retaining cells (LRCs) are CD45+. Scale bars, 20 μm. Quantifications of individual-keratinocyte H2BGFP intensity levels are presented underneath as histograms for each time point (in green). Best fits for the SP model with exponential- (grey) or gamma-distributed cell-cycle periods (orange lines) are shown. (C, E, G) Best estimates for the (gamma) distribution of the keratinocyte cell-cycle periods in hindpaw, ear and dorsal epidermis, respectively, as estimated from fits to each corresponding H2BGFP dilution data. Conservative solutions (in dark orange) are used for further inference. Vertical blue lines: average cell cycle period per site: <λ> = 2.0, 1.5, 1.2/week for paw (C), ear (E) and dorsum (G), respectively.

**Fig. S3.**
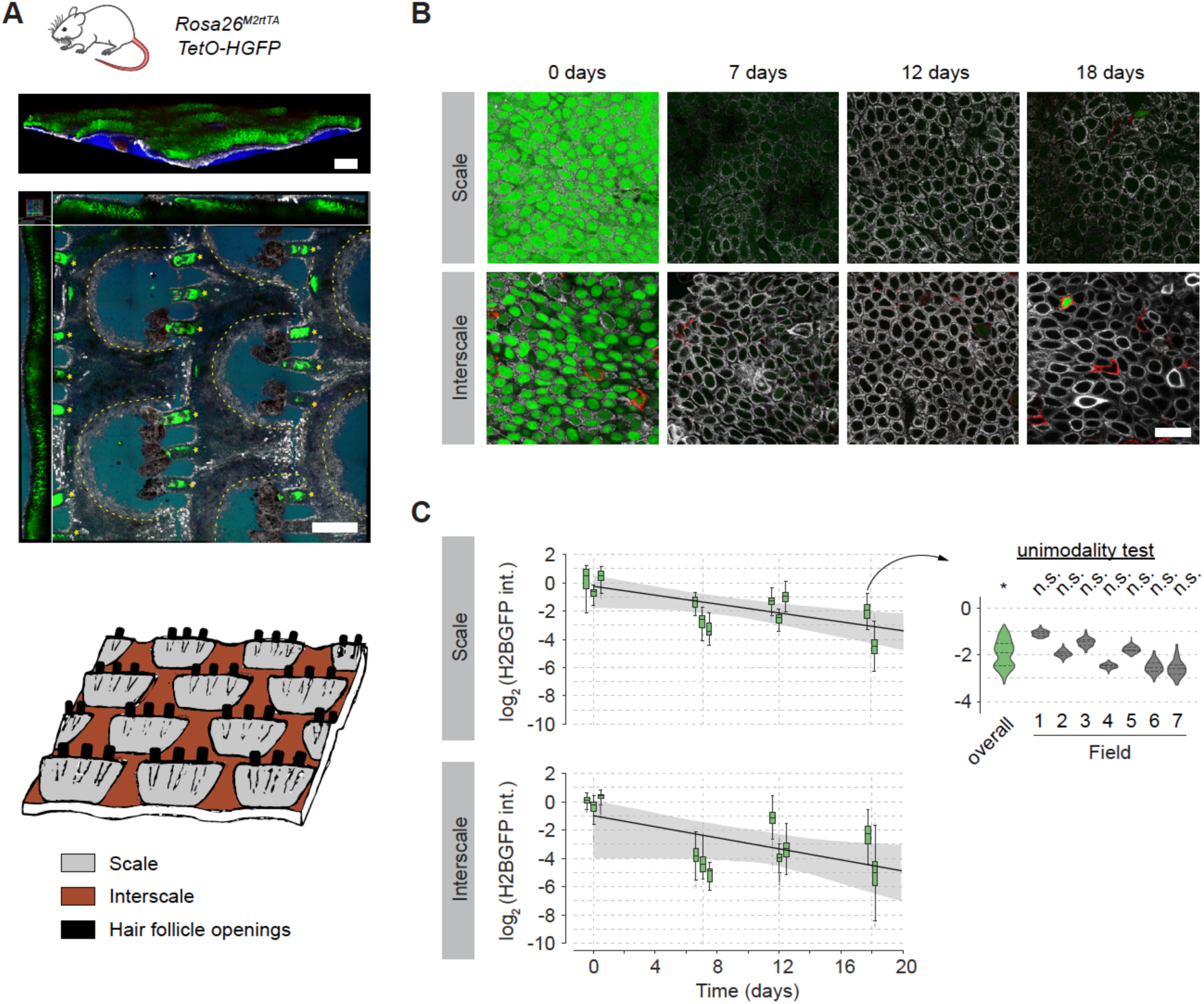
H2BGFP dilution analysis in tail epidermis. (A) Structure of the mouse tail epidermis. 3D reconstruction of confocal z stacks (top panel) and orthogonal xyz view (mid panel). Apical blister-shaped regions (scale) alternate with deeper regions (interscale) (boundaries delineated by dotted lines), being arranged in lines separated by hair follicles (asterisks). Green: H2BGFP expression; white: KRT14 immunostaining (as a marker of basal layer); red: CD45 immunostaining; blue: DAPI. Scale bars, 200 μm. Bottom panel: Cartoon illustrating the tail skin structure. (B) Representative confocal z stacks of scale and interscale regions of tail epidermis during H2BGFP chase experiments in *R26^M2rtTA^/TetO-H2BGFP* mice, showing H2BGFP (green), immunostaining for KRT14 (white) and immunostaining for pan-leukocyte marker CD45 (red). Scale bars, 20 μm. (C) Time course of basal-layer keratinocyte H2BGFP intensity distributions from scale and interscale regions of tail. Experimental data shown as boxplots per individual mouse (intensities normalized to average keratinocyte intensity at time 0). Solid black lines: average H2BGFP dilution rates (within 95% CI limits – shaded grey areas – where the value of λ=1.2/week reported by ^4^ falls). Insert: Detail of H2BGFP intensity distributions separated per field of view for the single tissue found to be bimodal in the overall, per-animal modality test (Fig. 2C). Analyses per field of view all resulted non-significant (unimodality).

**Fig. S4.**
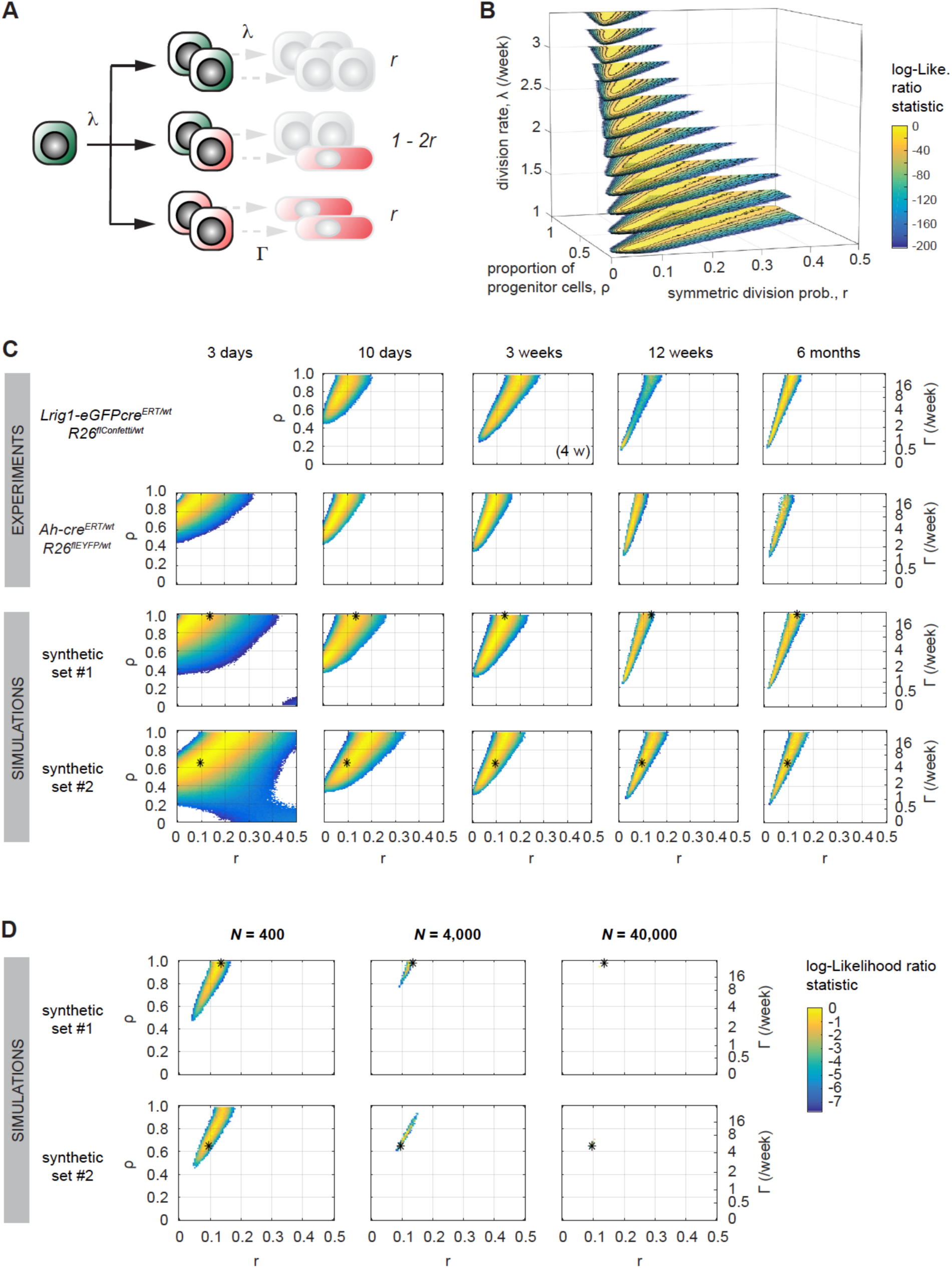
Single progenitor model parameter inference. (A) Parameterization of the SP model proposed to recapitulate the stochastic cell fate dynamics in epithelial homeostasis. In green, dividing cells; in red, differentiating cells (dim colors illustrate the possible lag in basal cell priming during its lifetime); in grey, unknown fate. λ: cell division rate. Γ: stratification rate. *r*: probability of sister cells undergoing the same fate (either both dividing or both differentiating) (the fraction of proliferating basal cells ρ is a dependent parameter, related to the value of Γ in homeostasis: ρ = Γ/(λ+Γ)). (B) MLE inference of the SP model parameters from experimental basal clone sizes. Data from lineage tracing in *Lrig1-eGFPcre^ERT/wt^ R26^flConfetti/wt^* mice esophagus are taken for illustration. 3D parameter solutions are color-coded according to likelihood value (fittings worsen as the log-likelihood ratio statistic gets more negative; values < −200 are not displayed). Without prior knowledge on cell proliferation rates, it is hard to resolve the dynamics, with semi-optimal solutions (yellow regions) spread across multiple values of λ. (C) 2D heat maps showing most likely parameter values obtained when analyzing, under the SP paradigm – with prior knowledge on λ – and for each time point separately, experimental *Lrig1*- and *Ah*-*Cre^ERT^* ^5^ derived clonal data from esophagus and, in parallel, synthetic clonal data simulated under specific parameter conditions (shown in asterisks) and subjected to reanalysis. In all cases, 100 randomly-chosen clones were analyzed per data set per time to correct for possible biases due to differences in sample size. Results are averages from bootstrapping. (D) SP model parameter discrimination improves as increasing the sample size. 100, 1,000 or 10,000 clones were sampled per time point from the synthetic data sets mentioned above, and the inference analyses repeated pooling the info from time points 1, 2, 4, and 6 weeks. Results shown as bootstrapping averages.

**Fig. S5.**
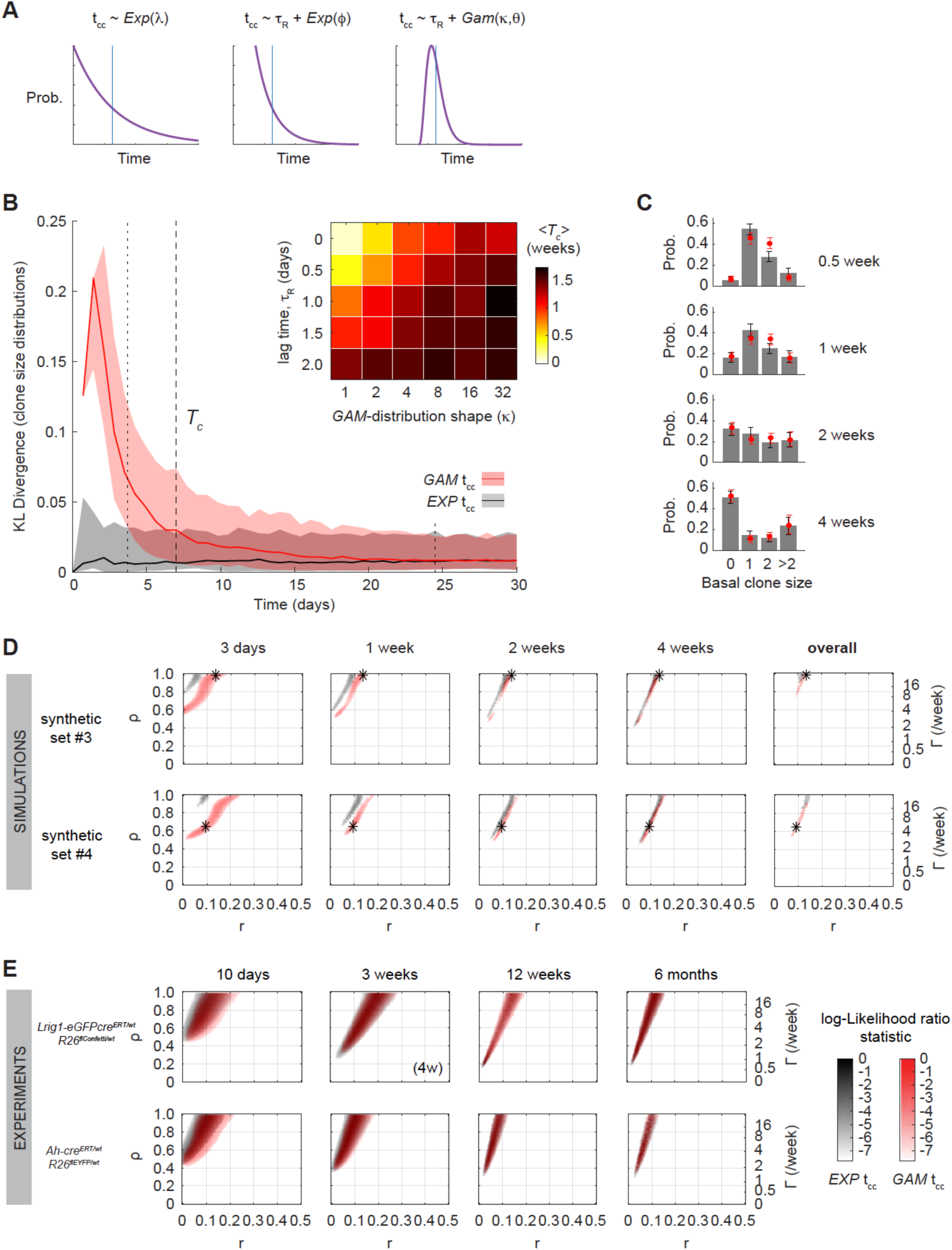
Assumptions on cell-cycle period distribution impact clone-dynamics parameter inference. (A) Possible models for the distribution of the individual cell-cycle period (t_cc_): exponential; delayed-exponential, where cells can only divide after a minimum refractory period τ_R_; delayed gamma distribution (with shape controlled by parameter κ). Blue vertical line: average cell-cycle length. (B) Time evolution of the Kullback-Leibler (KL) divergence between basal clone-size frequencies obtained from a limited set of simulated clones (300 clones per time point) following an exponential (grey) or gamma (red; τ_R_=0.5 days, κ=8) t_cc_, and the theoretical clone-size distribution predicted for an idealized infinite population under the exponential t_cc_ paradigm (lower values indicate a closer match). Solid lines and shaded regions stand for the average and 95% confidence intervals respectively on KL values from at least 100 independent sets. The average critical time *T_c_* (dashed line) after which clone-size distributions from gamma and exponential t_cc_ assumptions converge is reported as a heat map for various shapes of the cell-cycle distribution (all simulations carried out with λ=2.9/week, ρ=0.65, *r*=0.1). (C) Detail of differences in simulated basal clone-size distributions obtained at early time points under an exponential- (grey) vs. gamma- (red) distributed t_cc_ (parameter values as above). Data shown as average ± 95% CI on frequencies. (D) Parameter inference obtained at different time points from simulated clone-size datasets (1,000 clones per time point) run under the paradigm of a gamma-distributed t_cc_ (τ_R_=0.5 days, κ=8). In red scale: accepted parameter values when considering the actual t_cc_ distribution as a given prior. In grey scale are results assuming a default exponential t_cc_ distribution. Results are averages from bootstrapping. Neglecting the details of the cell-cycle period distribution can lead to deviations from the true parameter values (in asterisks). Similarly, parameter estimates obtained from *Lrig1-* and *Ah-Cre^ERT^* ^5^ derived experimental data sets in esophagus (100 clones per time point) do vary depending on the t_cc_ assumptions used in the inference analysis (E).

**Fig. S6.**
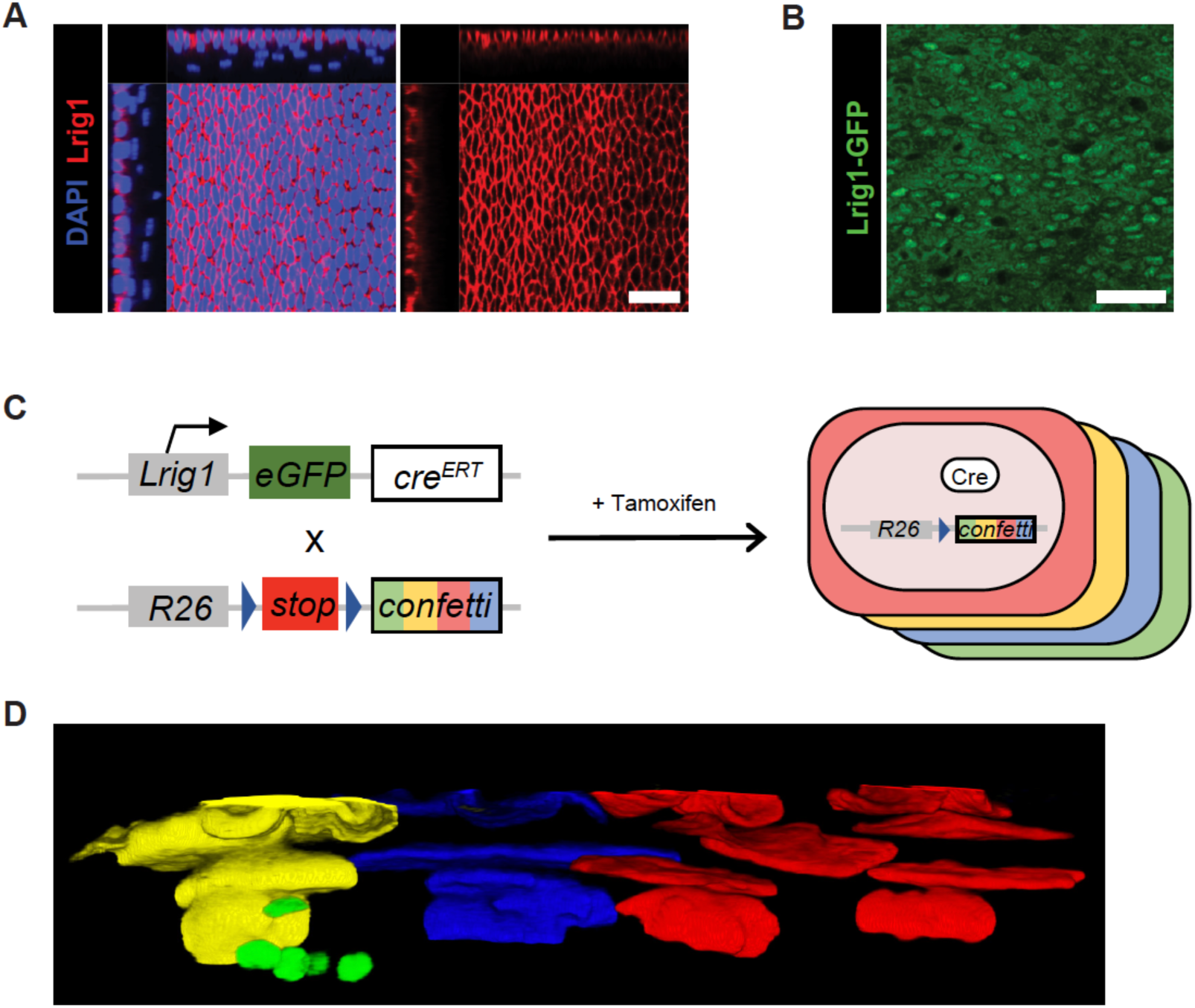
Experimental model used for lineage tracing in esophageal epithelium. (A) xyz planes view of confocal images showing *Lrig1* expression in wild type mice confined to the basal layer of the esophageal epithelium, where it is widespread. Red: LRIG1 immunostaining; blue: DAPI. Scale bar, 40 μm. (B) Rendered confocal z stacks of the basal layer in *Lrig1-eGFP-cre^ERT^* mice showing *Lrig1*-driven GFP expression. Scale bar, 40 μm. (C) Description of the transgenic mouse model used for lineage tracing. An *eGFP-IRES cre^ERT2^* construct was inserted in the exon 1 of the endogenous *Lrig1* locus, and a conditional confetti expression construct containing a “stop” cassette flanked by *LoxP* sites was targeted to the ubiquitous Rosa26 promoter. Upon induction with tamoxifen, Cre^ERT^ protein can migrate to the nuclei and excise the stop codon, resulting in the expression of one of the four different fluorescent proteins coded by confetti: GFP, YFP, RFP or CFP. Labelled, recombinant cells and their progeny (i.e. clones) can then be analyzed at different times. (D) 3D reconstruction of a confocal z stacks showing the side-view of an esophageal wholemount after induction in *Lrig1-eGFPcre^ERT/wt^ R26^flConfetti/wt^* animals.

**Fig. S7.**
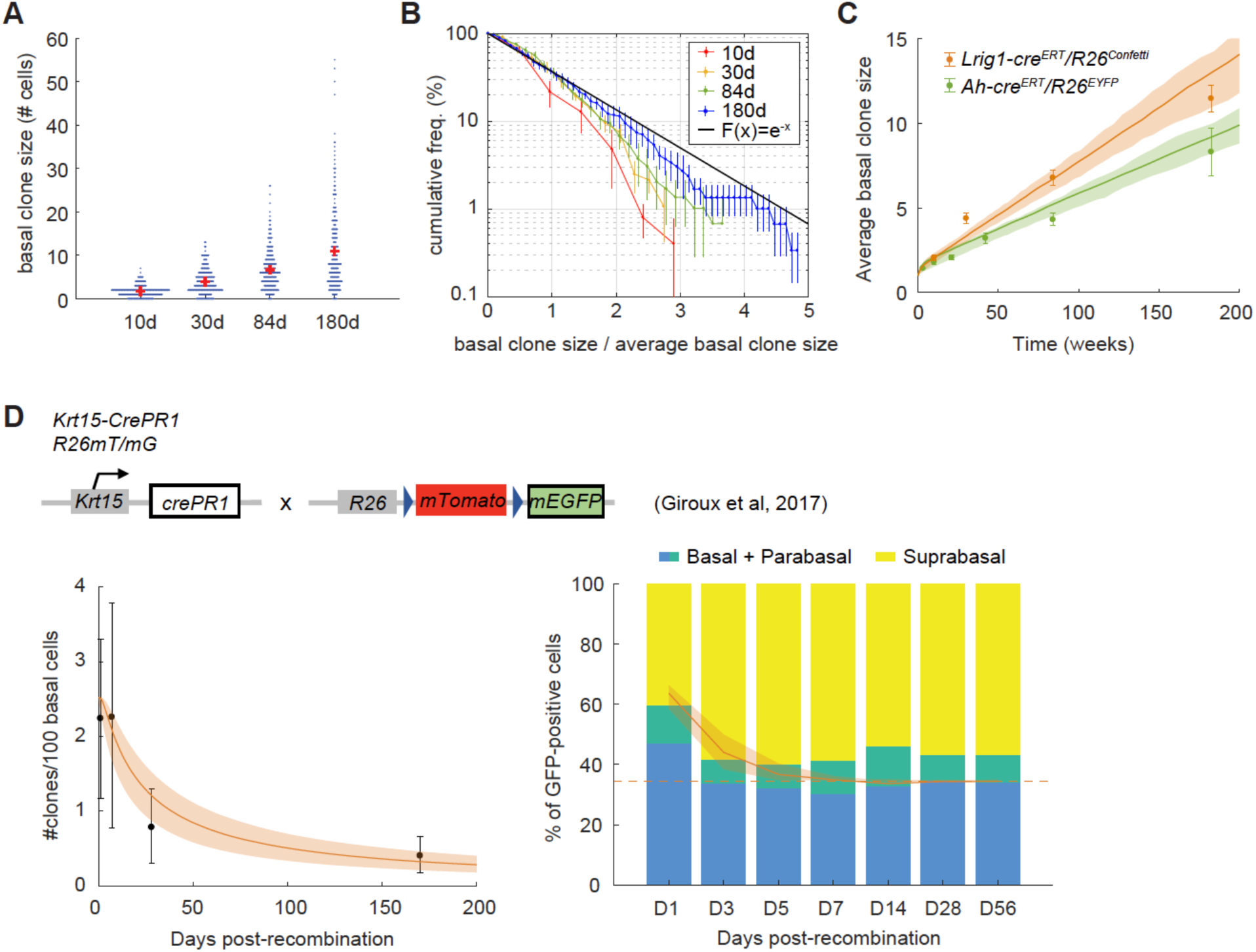
Labelled clone dynamics in Lrig1-cre^ERT^/R26^Confetti^ mouse esophagus and comparison to other lineage-tracing data sets. (A) Beeswarm plot showing Lrig1-derived basal-layer clone sizes (number of basal cells/clone) over time in the esophageal epithelium. Red marks show average size at each time. Only *surviving* clones, containing at least 1 basal cell, were considered. (B) Histogram of cumulative clone size frequencies normalized to the average basal clone size at each time (dots with error bars: mean ± s.e.m. from n=3 mice). At long term, distributions converge into a scaling behavior, where the probability of seeing clones of sizes larger than x times the average becomes constant and follows an exponential F(x) = e^-x^. (C) Time evolution of the average basal-layer size of surviving clones in induced *Ah-cre^ERT^ R26^EYFP^* mice ^5^ as compared to *Lrig1-eGFP-cre^ERT^ R26^flConfetti^*. Experimental data shown as dots with errorbars (mean ± SEM from n ≥ 3 animals). SP model fits shown in green and orange, respectively (solid lines stand for the MLE; light areas: 95% CI). (D) The inferred keratinocyte cell behavior from the *Lrig1-eGFP-cre^ERT^ R26^flConfetti^* system fits independent experimental lineage-tracing data from *Krt15-cre^PR1^ R26^mT/mG^* mouse esophagus ^29^, reproducing the decay in the basal clone density over time (left panel) and the change in the proportion of labelled cells located in basal and suprabasal compartments (right panel). Experimental sets are from Fig. S3 and Fig. 2 in ^29^, respectively. Model fits shown in orange (solid lines: MLE; light areas: 95% CI; dashed line corresponds with the estimated fraction of basal cells in homeostasis).

**Fig. S8.**
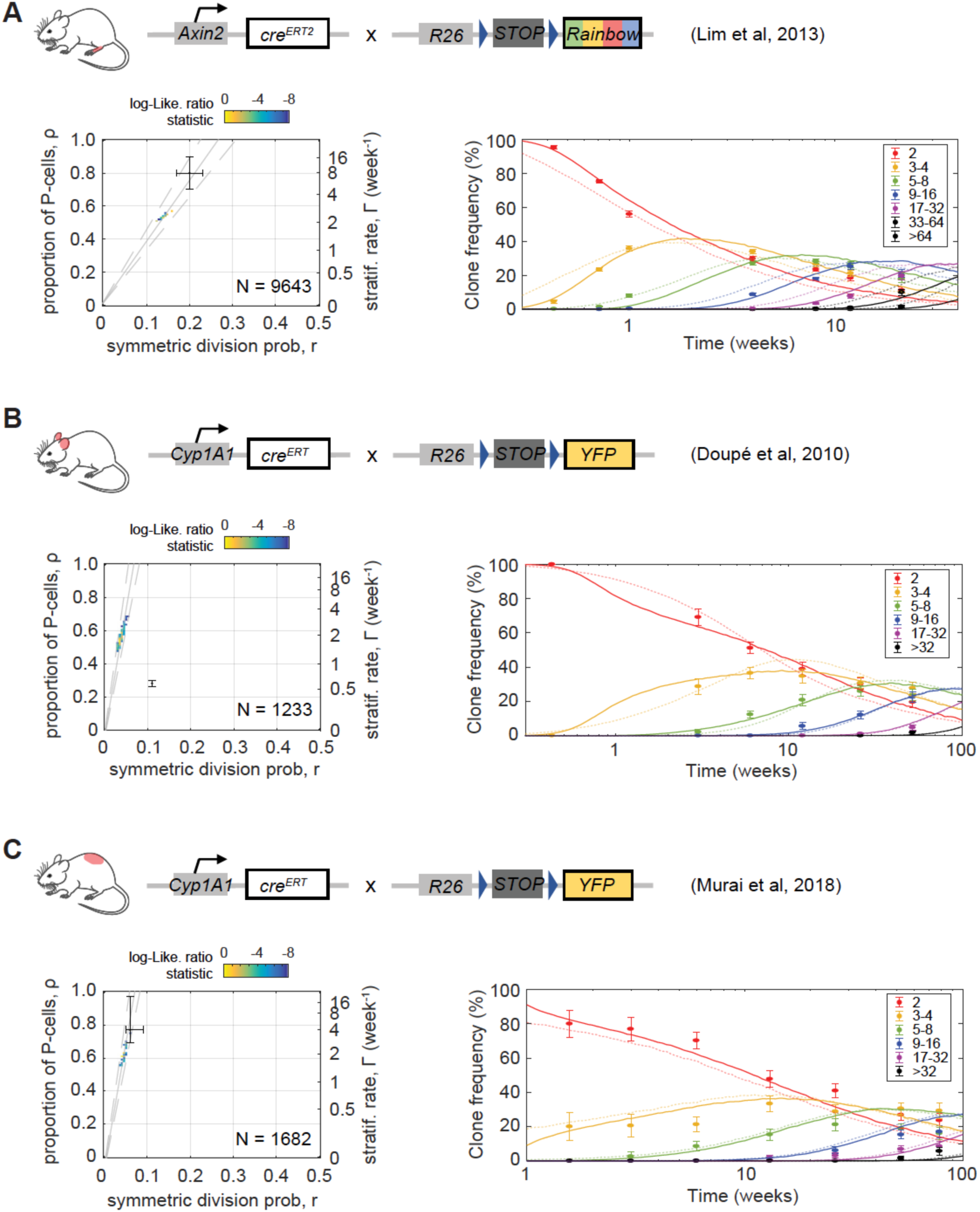
The single-progenitor model fits clone dynamics in different regions of skin epidermis. (A, B, C) left panels: SP model parameter inference on lineage tracing datasets from paw epidermis ^11^ using *Axin2-cre^ERT^R26^Rainbow^* animals (A), and ear ^30^ and dorsal ^21^ interfollicular epidermis in *AhYFP* mice (B and C, respectively). Parameter estimates are obtained by MLE based on SP model simulations constrained by the cell-cycle period distribution inferred from each corresponding cell proliferation analysis. Regions within the dashed grey lines fall consistent with the predicted ρ/*r* ratios from the linear scaling of the average clone size. The total number of clones counted in each data set is displayed in the corresponding graph and previous parameter estimates given in the original publications shown as black error bars. (A, B, C) right panels: Experimental *Axin2-* (A; from ^11^) and *Ah*- (B and C; from ^30^ and ^21^) derived basal-layer clone sizes (dots with error bars as standard error of proportion) are excellently fit with the SP model with gamma-distributed cell cycle times (lines; prediction from MLE). Dim dashed lines: fits obtained with parameter estimates given in the original publications. Frequencies for each clone size (basal cell number) are shown in different colors.

**Fig. S9.**
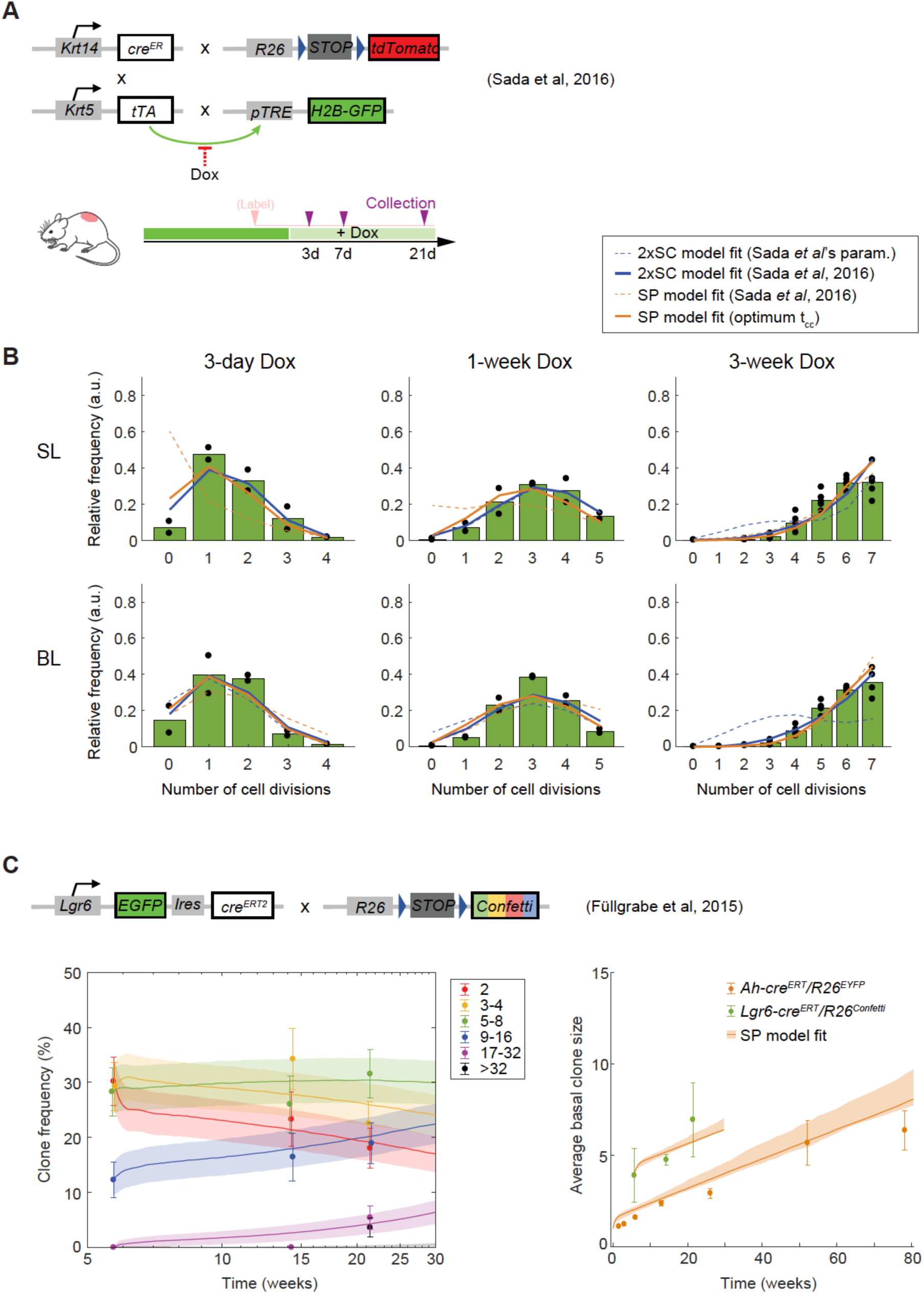
The single-progenitor model fits independent transgenic-mouse data sets on keratinocyte behavior in dorsal skin. (A) Transgenic mouse model used by ^19^ to study epidermal cell proliferation in dorsal skin. Unlike *R26^M2rtTA^/TetO-H2BGFP* mice (Tet-On), *Krt5^tTA^/pTRE-H2BGFP* mice show constitutive H2BGFP expression, and treatment with Dox is required for repression of H2BGFP production during the duration of the dilution experiment (Tet-Off; see Fig. S1). (B) Reanalysis of H2BGFP dilution data in ^19^. Distributions of cell division number after different chase periods, as deconvoluted from H2BGFP fluorescence histograms of FACS sorted basal and suprabasal (spinous-layer) cells (green bars; dots: individual-mice data). Data extracted from Fig. 3 in ^19^. Our computational-simulation fits using a SP model with gamma-distributed cell-cycle times (solid orange lines) mimic the best fits provided by ^19^ using a 2xSC model (solid blue lines), while the latter results in deviated fits with the parameters claimed in their text (dashed blue lines). Poor SP-model fits from ^19^ (dashed orange lines) were due to oversimplified exponential cell-cycle assumptions. (C) The inferred keratinocyte cell behavior from *AhYFP* mouse back skin epidermis fits independent experimental lineage-tracing data from *Lgr6-eGFPcre^ERT^ R26^flConfetti^* mice ^33^. Left panel: Empirical long-term, basal-layer clone size distributions are displayed as mean frequency ± standard error of proportion (dots with error bars) for each clone size or basal cell number (in different colors). Lines and shaded areas correspond to the SP-model MLE predictions and ranges within ± SD from bootstrapping (random sampling of experimental clone sizes at P8w were considered as starting condition for simulations on the *Lgr6*-based system, to account for the fact that induction occurred early in development, before mouse epithelia become homeostatic). Right panel: Time evolution of the average basal-layer size of surviving clones in *Lgr6-eGFPcre^ERT^ R26^flConfetti^* mice (from ^33^) as compared to *Ah-cre^ERT^ R26^EYFP^* mice ^21^. Experimental data shown as dots with error bars (mean ± s.e.m. from n ≥ 3 animals). SP model fits shown in orange (solid lines stand for the MLE; light areas: 95% CI), matching both average trends from single-labelled progenitors in the *Ah*-based data set and preformed clones in the *Lgr6*-based system.

**Fig. S10.**
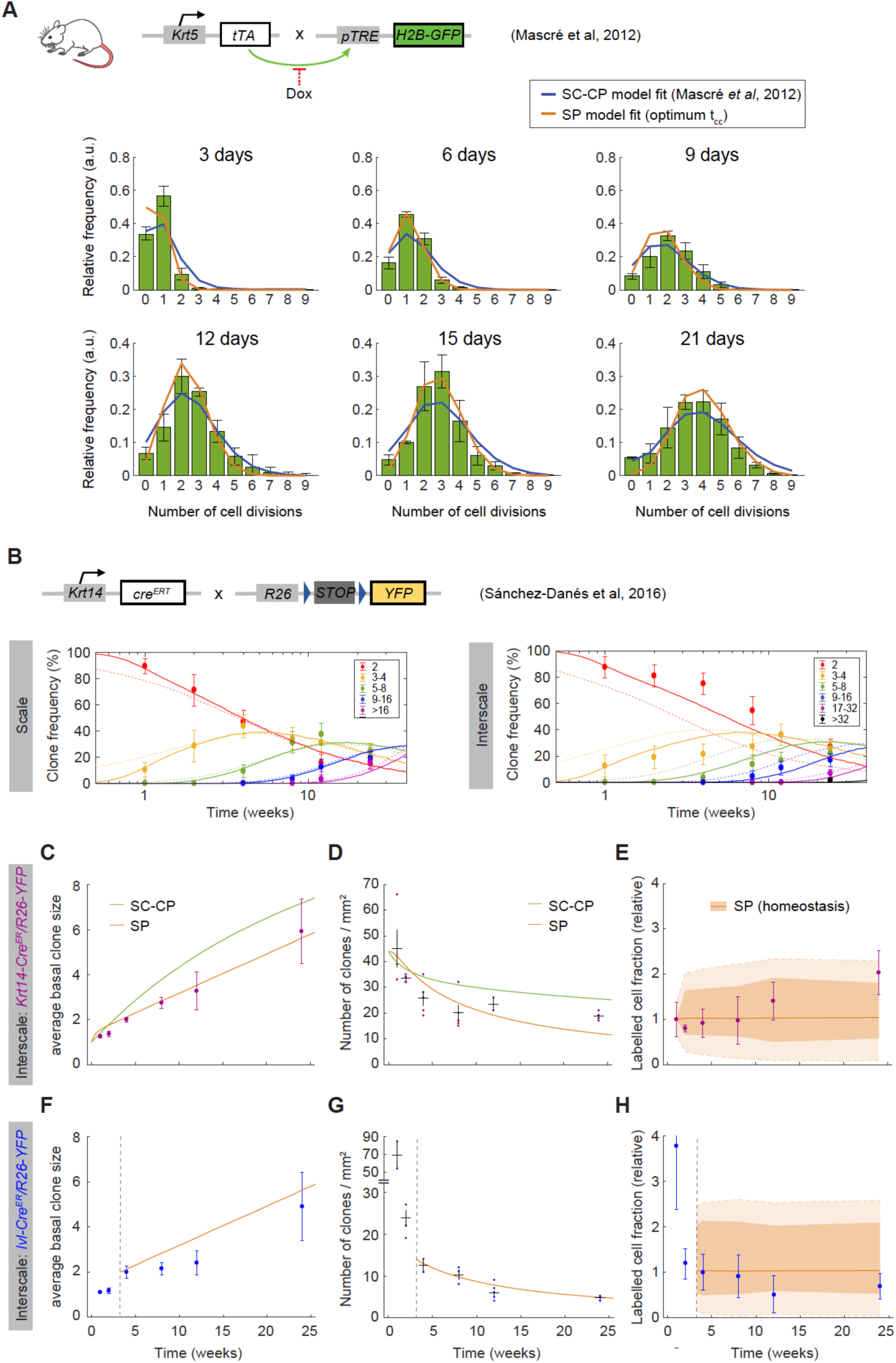
The single-progenitor model fits different transgenic-mouse data sets on keratinocyte behavior in tail skin. (A) Reanalysis of H2BGFP dilution data from *Krt5^tTA^/pTRE-H2BGFP* (Tet-Off; see Fig. S1) mouse tail skin ^4^. Distributions of cell division number after different chase periods, as deconvoluted from H2BGFP fluorescence histograms of FACS sorted basal cells (green bars with error bars: mean ± s.e.m. from n = 3 animals). Data extracted from Fig. 3 in ^4^. Our computational simulations using a SP model with gamma-distributed cell-cycle times (solid orange lines) show as good a fit to the data as the more complex SC-CP model used by ^4^ (solid blue lines). Unrealistic, exponential cell-cycle time distributions were considered in the original publication. (B) Basal clone size distributions in scale and interscale regions of tail following lineage tracing in *Krt14-cre^ERT^ R26^YFP^* mice ^20^. Frequencies for each clone size are in different colors (dots with error bars: empirical values reported, along with standard error of the proportion). Solid lines: SP-model MLE fits obtained using as a prior the value for the average division rate measured by ^4^ and a reasonable gamma-shaped cell-cycle time distribution. Dim dashed lines: fits obtained using the parameterized models described in ^20^, where a SC-CP scenario is considered in interscale. (C, D, F, G) Time course of the average basal-layer size and density of clones labelled with *Cre^ERT^* expressed from *Krt14* or *Ivl* promoters in interscale (purple and blue datasets, respectively; dots with error bars: mean and SEM from n ≥ 2 mice) ^20^. Orange lines: fits obtained with the SP model with same parameter values determined from (B). Green lines: fits on *Krt14*-derived data obtained from the parameterized SC-CP model used by ^20^. (E, H) Changes in the labelled cell fraction derived for both *Krt14-cre^ERT^ R26^YFP^* (E) and *Ivl-cre^ERT^ R26^YFP^* (H) experimental model data sets (error bars: mean ± s.d.) (inferred by multiplying average labelled clone densities by average clone sizes at each time, accounting for error propagation). Possible experimental trends fall within the domain of uncertainty given by the SP model in homeostasis considering the large errors due to the actual limited sample size (dark orange area: 95% CI on the mean; light orange area: margins given with ± s.d.) (see Supplementary Theory for details).

