## Supplementary Methods for "The single-progenitor model as the unifying paradigm of squamous epithelial maintenance"

### Supplementary Methods: Theory

This Supplementary Methods report is intended to provide a detailed description of the methodology used in the Main Text and the quantitative arguments supporting the paradigm of a single population of progenitor cells in squamous epithelia. In **section 1** we describe some control measures to test the adequacy of our experimental system for lineage tracing in esophagus. In **section 2** we formulate the different stochastic models of cell behavior and discuss the limitations that clonal dynamics features present for model discrimination. In **section 3** we describe the methods used to infer a single mode of keratinocyte cell proliferation. **Section 4** follows with the combined approach used for model inference on clonal lineage-tracing data sets constrained by cell-cycle properties. Finally, in **section 5** we revisit the quantitative methodology and arguments from previous publications.

#### 1. Lineage-tracing controls and labelling representativeness

In the main text, we describe a new lineage tracing experiment in esophageal epithelium using inducible *Lrig1-eGFPcre*$^{ERT}$ *R26*$^{flConfetti}$ mice. Before introducing the quantitative methods and theory involved in clonal fate analysis, in this section we address some controls regarding the labelling representativeness of self-renewal in this tissue.

*Lrig1* was ubiquitously expressed throughout the basal layer of the esophageal epithelium, where proliferation is confined (immunostaining; **Fig. S6A**), and *Lrig1*-driven GFP expression was detectable in more than 94% of the basal cells (**Fig. S6B**). This argues that *Lrig1*-derived labelled clones widely represent basal keratinocyte dynamics in the esophageal epithelium.

An inducible *Cre*/*Confetti* reporter was used to track the fate of *Lrig1*-expressing basal cells (**Fig. S6C**). To exclude the possibility of spontaneous florescence reporter expression, esophageal epithelia were collected from uninduced, 12-16 week old mice ($N$=3), confirming the absence of any labelled clone and hence of any leakage (**Table S5**). Following low-dose tamoxifen administration, CFP-, RFP- and YFP-labelled cell patches were analyzed for clonal behavior at different times post-induction. The low total labelling efficiency (1 in 301 ± 106 (mean ± SEM) basal cells by 10 days post induction; **Table S5**) and distinction of different fluorescent colors minimizes the risk of clonal merging [1], so that single-color patches were considered clonal. Induced GFP-labelled cells were excluded from the analysis given the expression of GFP from the *Lrig1* locus.

The number of basal cells per clone, $n$ (basal clone size), was counted, and the frequency of clones of a certain size $x_n$ reported for each individual label reporter and time (**Table S5**). No statistical differences in the distributions of clone size frequencies were seen between CFP, RFP and YFP labelled cell populations at any given time (Kruskal-Wallis test, p = 0.17, 0.27, 0.99, 0.22 at time 10d, 30d, 84d, 180d, respectively; non-significant too by pairwise comparisons using Kolmogorov-Smirnov test), justifying pooling the data from the different label reporters. Just in the case where the time courses in the number of clones per unit area and the proportion of labelled basal cells were shown, only RFP clones were considered, given the low, variable induction of the other florescent reporters (including these numbers did not alter the conclusions).

Importantly, overall, the fraction of Confetti labelled basal cells remained approximately constant over time (**Fig. 3C**). This agrees with homeostatic behavior and strongly supports the hypothesis that the labelled cell population is representative of the epithelial self-renewal.

#### 2. The possible models of epithelial cell dynamics

Homeostasis is a fundamental feature of adult tissues. Whilst the specific mechanisms for homeostasis may vary depending on the organization of the niche itself, within squamous epithelia several models have been proposed that are sufficient to explain tissue maintenance. These can be broadly separated based on two key features: the presence of a single or multiple dividing population, and whether the progeny fates are balanced by a deterministic or stochastic process. Deterministic processes (*invariant asymmetric self-renewal*) lead to the growth of stable, similar sized clones over long periods of time following labelling of stem cells, regardless of the presence or absence of transient amplifying cells (cells that have a limited division capacity) [2]. In contrast, stochastic models (*population asymmetric self-renewal*) achieve balance by allowing individual cells to divide either asymmetrically or symmetrically, with an equal probability of symmetric stem or differentiation fates (**Fig. 1D**). A consequence of these branching processes is that clones exist in neutral competition, and develop heterogeneous sizes over time as some grow, whilst others diminish or even become extinct. Here we briefly review this latter class of models and the evidence that supports them.

Several features of the clone size distributions arise from population asymmetric self-renewal [1, 3]:

- The number of surviving labelled clones, containing at least one basal cell, continuously decays over time, following a hyperbolic function, $P_{surv}(t) \sim 1/t$. This reflects the non-null probability of clone extinction.
- The average number of basal cells in the surviving clones grows linearly with time, $\langle n \rangle_{surv}(t) \sim t$, to compensate the decline in clone density, so that the overall cell population remains constant.
- The surviving clones display increasingly heterogeneous sizes. The distribution of basal clone sizes adopts a scaling behavior at long term, so that the chance of finding a clone larger than some multiple of the average becomes constant, i.e. $P_n^{cum}(t) = f[n/\langle n(t) \rangle_{surv}]$, where, typically, $f(x) = e^{-x}$.

We found that these hallmarks are all fulfilled by the *Lrig1-eGFPcre^ERT R26^flConfetti* clonal data (**Fig. 3**; **Fig. S7A,B**), and also shared by lineage-tracing datasets across different skin territories (**Fig. S8**), confirming self-renewal dynamics in mouse squamous epithelia is dominated by stochastic cell fates comprising both symmetric and asymmetric division outcomes.

However, it remains contested whether this stochastic clone dynamics is underpinned by a single population of dividing cells [3] or multiple populations. In different studies these multiple populations have been proposed either to coexist independently within a tissue [4] or with a hierarchical relationship between them [5] (**Fig. 1D**). In the following subsections we formally describe the different models and explain their specific characteristics.

*The single-progenitor (SP) model:*

In the *single-progenitor model*, there is a single population of progenitor cells (P cells), which divide regularly, with an average rate λ, to give rise, with a certain probability, to either two daughter progenitors (P + P), two differentiating cells (D + D) or one of each (P + D) [1] (**Fig.

**1D; Fig. S4A**). Differentiating cells (D) are post-mitotic and leave the basal layer with stratification rate Γ, constituting suprabasal-layer cells that are ultimately shed[1].

$$P \xrightarrow{\lambda} \begin{cases} P+P & \text{Prob. } r \\ P+D & \text{Prob. } 1-2r \\ D+D & \text{Prob. } r \end{cases} \qquad (1)$$
$$D \xrightarrow{\Gamma} \emptyset$$

In order to ensure tissue homeostasis, the probabilities of symmetric divisions that lead to two proliferating cells or two differentiated cells are balanced, i.e. both are defined by a fixed parameter $r \in [0, 0.5]$. This sets a total of three unknown parameters $\theta = \{\lambda, r, \Gamma\}$. Furthermore, under these homeostatic conditions, one can assume that the proportion of progenitor cells in the basal layer, denoted as ρ, remains constant, and overall, the net rate at which post-mitotic cells are generated in the basal layer gets compensated by cell stratification (and shedding), so that $\rho = \Gamma / (\Gamma + \lambda)$. It follows that unless the stratification rate is huge ($\Gamma >> \lambda$), the basal compartment would still show a substantial level of heterogeneity (with a mixture of both P and D cell pools).

*The two independent stem-cell (2xSC) model:*

In this alternative model, derived from Sada et al [4] (see original formulation in **section 5**), two independent stochastic proliferating populations of stem cells are considered (2xSC model)[4]. These $S_1$ and $S_2$ populations divide at two different rates, $\lambda_{S1} << \lambda_{S2}$, and each follows a pattern of stochastic fate choices similar as in Eq. (1) with a given probability of symmetric division outcome, $r_{S1}$ and $r_{S2}$, allowing duplication or differentiation. We make the simplifying assumption that $r_{S1} = r_{S2} = r$.

$$S_1 \xrightarrow{\lambda_{S1}} \begin{cases} S_1+S_1 & \text{Prob. } r \\ S_1+D & \text{Prob. } 1-2r \\ D+D & \text{Prob. } r \end{cases}$$

$$S_2 \xrightarrow{\lambda_{S2}} \begin{cases} S_2+S_2 & \text{Prob. } r \\ S_2+D & \text{Prob. } 1-2r \\ D+D & \text{Prob. } r \end{cases} \qquad (2)$$
$$D \xrightarrow{\Gamma} \emptyset$$

This model yields five adjustable parameters $\theta = \{\lambda_{S1}, \lambda_{S2}, r, \Gamma, \rho_{S1}\}$, where $\rho_{S1}$ is the fraction of slow-dividing stem cells in the basal layer. One can retrieve the proportion of fast-cycling stem cells $\rho_{S2}$, or the proportion of differentiating cells $\rho_D = 1 - \rho_{S1} - \rho_{S2}$, from the parameter ratios given by the condition of homeostasis. In this way, $\rho_{S2} = (\Gamma - \rho_{S1}(\Gamma + \lambda_{S1})) / (\Gamma + \lambda_{S2})$. Alternatively, we can express the relative fraction of proliferating cells that are slow-cycling stem cells in homeostasis, $\chi_{S1}^{div}$:

$$\chi_{S1}^{div} = \frac{\rho_{S1}(\lambda_{S2}+\Gamma)}{\rho_{S1}(\lambda_{S2}-\lambda_{S1})+\Gamma} \qquad (3)$$

*The hierarchical stem cell/committed progenitor (SC-CP) model:*

---

[1] We have omitted the dynamics in the suprabasal compartment as these cells do not contribute to tissue maintenance.

In this model, introduced by [5] and revisited in [6], a slow-cycling population of stem cells is considered to underpin the self-renewing dynamics of a second, quickly-dividing population of progenitor cells[6]. Stem (S) cells divide at a slow rate $\lambda_S$ and undergo stochastic fates, so that they generate, with a certain probability, either two daughter stem cells (S + S), two progenitor cells (P + P) or one of each (S + P). Progenitor cells in turn divide at a faster rate, $\lambda_P >> \lambda_S$, and commit to stochastic fates too upon division, giving rise to progenitors or differentiating cells as previously indicated. Mascré et al [5]made the assumption that $r_S = r_P = r$ (depicted below). In contrast, Sánchez-Danés et al [6] assumed these probabilities varied independently.

$$
\begin{aligned}
&S \xrightarrow{\lambda_S} \begin{cases} S+S & \text{Prob. } r \\ S+P & \text{Prob. } 1-2r \\ P+P & \text{Prob. } r \end{cases} \\
&P \xrightarrow{\lambda_P} \begin{cases} P+P & \text{Prob. } r(1-\Delta) \\ P+D & \text{Prob. } 1-2r \\ D+D & \text{Prob. } r(1+\Delta) \end{cases} \\
&D \xrightarrow{\Gamma} \emptyset
\end{aligned} \tag{4}
$$

The hierarchical proliferative structure makes necessary to introduce a probabilistic bias ($\Delta \in [0,1]$) in the progenitor daughter fates, so that on average progenitor cells differentiate more often than what they duplicate to compensate for their net production from the stem cell pool and thus guarantee tissue homeostasis. As a result, there are five unknown parameters $\theta = \{\lambda_S, \lambda_P, r, \Gamma, \Delta\}$. The following relationships between parameters can be established at homeostasis:

$$
\begin{aligned}
&\lambda_S S^{ss} = 2\lambda_p \Delta r_P P^{ss} \\
&\lambda_P P^{ss}(1 + 2\Delta r_P) = \Gamma D^{ss}
\end{aligned} \tag{5}
$$

where $S^{ss}$, $P^{ss}$, and $D^{ss}$ represent the bulk populations of stem cells, progenitors and differentiating cells in the basal layer at homeostasis, satisfying the stationary-state conditions d$S$/dt = 0, d$P$/dt = 0, and d$D$/dt = 0, respectively. Rearranging Eq. 5, we get:

$$
\begin{aligned}
&\rho_S = \frac{\omega/\lambda_S}{\omega/\lambda_S + 1 + \lambda_P/\Gamma + \omega/\Gamma} \\
&\rho_P = \frac{1}{\omega/\lambda_S + 1 + \lambda_P/\Gamma + \omega/\Gamma} \\
&\rho_D = \frac{\lambda_P/\Gamma + \omega/\Gamma}{\omega/\lambda_S + 1 + \lambda_P/\Gamma + \omega/\Gamma}
\end{aligned} \tag{6}
$$

where $\rho_x$ represent the proportion of each x-cell type in the basal layer, and $\omega = 2\lambda_p \Delta r_P$. From here one could deduce the relative fraction of proliferating cells that would represent slow-cycling stem cells in homeostasis, i.e. $\chi_S^{div}$:

$$
\chi_S^{div} = \frac{S^{ss}}{S^{ss} + P^{ss}} = \rho_S/(\rho_S + \rho_P) = \frac{\omega}{\omega + \lambda_S} \tag{7}
$$

*Simulating stochastic clone dynamics under the different hypotheses:*

In order to explore the range of possible clone dynamics that these different models can offer, we formulated each in terms of its corresponding stochastic Master equation. For instance, for the SP model we have:

$$\frac{\partial P_{n_P,n_D}}{\partial t} = \lambda\left[r(n_P - 1)P_{n_P-1,n_D} + r(n_P + 1)P_{n_P+1,n_D-2} + (1 - 2r)n_P P_{n_P,n_D-1}\right] + \Gamma(n_D + 1)P_{n_P,n_D+1} - \lambda n_P P_{n_P,n_D} - \Gamma n_D P_{n_P,n_D} \quad (8)$$

where $\partial P_{n_P,n_D}/\partial t$ describes the time evolution of the probability of finding clones containing $n_P$ progenitor cells and $n_D$ differentiated cells. Due to the difficulty in computing the analytical solutions for these equations [7], in our analyses $P_n(t)$, the probability of each given basal clone size $n$, was estimated for each model from multiple simulations ($N$ = 100,000) of the Master equation, following Gillespie's algorithm by default [8, 9] (for more elaborated methods, see **sections 3,4**).

As initial condition, we generally set (except when stated otherwise) to start from a random, single labelled proliferative cell, since we can assume that any initially induced differentiating cell will be rapidly swept into the suprabasal compartment and therefore make a negligible contribution to the basal clone dynamics at medium-long term ($t > 1/\Gamma$). In this way, for the SP model: $P_n(0) = \delta_{n_P,1}\delta_{n_D,0}$, where $\delta_{n,m}$ represents the Kronecker delta. For the 2xSC model: $P_n(0) = \delta_{n_{S1},B(1,\chi_{S1}^{div})}\delta_{n_{S2},B(1,1-\chi_{S1}^{div})}\delta_{n_D,0}$; and for the SC-CP model: $P_n(0) = \delta_{n_S,B(1,\chi_S^{div})}\delta_{n_P,B(1,1-\chi_S^{div})}\delta_{n_D,0}$. Note that $B(m,p)$ represents a random binomial probability term in the two-dividing population models, so that on average, a fraction $\chi_{S1}^{div}$ of simulations initiate with a labelled cell targeting the slow-cycling population, and 1-$\chi_{S1}^{div}$ with a quickly-dividing cell, in proportions consistent with the actual ratio of these cell types in homeostasis (Eq. 3 and 7).

All three models show highly similar scaling behaviour. Regardless of the parameter values chosen, clonal dynamics under the SP model adopts the general scaling properties described earlier. In particular, at $t > 1/r\lambda$ the system enters an asymptotic regime [3] where:

$$\begin{aligned} P_{surv}(t) &= \frac{1}{1+r\lambda t} \\ \langle n\rangle_{surv}(t) &= \frac{1}{\rho} + \frac{r\lambda}{\rho}t \\ P_n^{cum}(t) &= \exp[-n/\langle n\rangle_{surv}(t)] \end{aligned} \quad (9)$$

The exponential scaling of the cumulative clone size frequencies yields a linear trend when representing $\log P_n^{cum}(t)$ vs. $n/\langle n\rangle_{surv}(t)$, as observed in oesophageal epithelium (**Fig. S7B**).

In the 2xSC and SC-CP models, the distribution of basal clone sizes converges to a shape where $\log P_n^{cum}(t)$ does not change fully linearly with the normalized basal clone sizes $n/\langle n\rangle_{surv}(t)$ but displays a biphasic pattern with a U-shaped curve for small clone frequencies due to the mixture of the two different proliferating populations with distinct potential to yield larger clone sizes. However, any deviation from the exponential scaling becomes negligible under most parameter conditions. Similarly, the average clone size and the fraction of surviving clones of these models adopt curved shapes that show only slight divergence from the single progenitor model for many parameter sets.

Finally, it is worth commenting on the predictions for the evolution of the labelled cell fraction. If the labelled cell fraction faithfully represents the proportions of proliferative cell-types in the homeostatic tissue and an adequate number of clone simulations were initiated from each subpopulation (starting with a single cell, in the way previously stated), the overall population of tracked (labelled) dividing cells across the 100,000 simulations $\pi^{div}$ remained approximately constant over time, regardless of the model considered. However, these dynamics could become sub-linear or supra-linear under the SC-CP hypothesis insofar as different initial ratios of S and P cells, $\chi_{S,label}^{div}$ and $1-\chi_{S,label}^{div}$, were tracked than those expressed in Eq. 7. This would reflect a scenario where labelling preferentially targets P cells or S cells, respectively. The time evolution of the labelled cell fraction (omitting the contribution of D cells) would be described by: $d\pi^{div}/Ndt = dS/dt + dP/dt$. Integrating, we get:

$$\pi^{div}(t) = \chi_{S,label}^{div}\left(1+\frac{\lambda_S}{\omega}(1-e^{-\omega t})\right) + \left(1-\chi_{S,label}^{div}\right)e^{-\omega t} \qquad (10)$$

This has motivated claims arguing that any deviation from a constant value over time can be attributed to imbalanced fate choices (SC-CP model dynamics, where one could primarily target one or the other dividing subpopulation) [6]. Nevertheless, we note that if we considered a non-negligible fraction of differentiating cells in the basal layer and the possibility of labelling these at a more or less extent, sub-linear or supra-linear trends in the labelled cell fraction could be expected too under the SP model paradigm, if $\rho_{P,label}$ and thus $\rho_{D,label} \equiv 1-\rho_{P,label}$ were different than the proportions given by $\rho$ in homeostasis. In this case, $d\pi/Ndt = dP/dt + dD/dt$. Integrating:

$$\pi(t) = \rho_{P,label}\left(1+\frac{\lambda_S}{\Gamma}(1-e^{-\Gamma t})\right) + \left(1-\rho_{P,label}\right)e^{-\Gamma t} \qquad (11)$$

Notice the similarity of this expression with that in Eq. 10, which relegates the point to a matter of differences in the time scales of the pre-asymptotic behaviour before settling to a constant value.

Altogether, these theoretical modelling results suggest that lineage tracing alone would provide little evidence to support one or another stochastic cell fate model.

#### 3. H2BGFP dilution analysis and inference on homogeneous keratinocyte cell behavior

Since the 2xSC and SC-CP models involve the existence of subpopulations of proliferating cells dividing at different rates, we speculated that we could discriminate between these scenarios and that of the SP model by analyzing the heterogeneity in the pattern of H2BGFP expression of individual keratinocytes during the time course of H2BGFP dilution experiments in transgenic *$R26^{M2rtTA}$/TetO-H2BGFP* mice. In this section we describe the experimental details and quantitative modelling involved in this analysis.

Cohorts of at least 2-3 *$R26^{M2rtTA}$/TetO-H2BGFP* mice were culled at 0, 7, 12 and 18 days post-doxycycline administration, and the epithelial basal-layer plane imaged from wholemounts of esophagus and hindpaw, ear and tail skin (**Fig. 2A**). At least 5-8 random fields of view were analyzed per tissue per animal, acquired from distant regions of epithelium. This was to guarantee as much as possible the tissue representativeness and control for possible regio-specific differences in the cellular turnover. Samples from back skin epidermis were acquired independently at 0, 5, 11, 14 and 21 days post-doxycycline treatment [10]. All images were processed with ImageJ to segment nuclear areas (based on DAPI staining) and quantify H2BGFP intensity levels in individual basal keratinocyte nuclei (intensity values were averaged over each nuclear area) (**Table S2**). Mitotic cells and $CD45^+$ (immune) cells were scored but excluded from the quantitative analysis. Also, immunostaining for Krt14 (a basal keratinocyte marker) helped to exclude suprabasal cells or other cell types from further analysis in tail skin epidermis, which is particularly wavy (**Fig. S3A**).

*Patterns of keratinocyte H2BGFP intensity distributions*

If all proliferating keratinocytes behaved as an equivalent population of progenitors dividing at a similar constant rate λ, we would expect a monotonous H2BGFP dilution pattern over time, where all individual-cell fluorescence intensities I(t) would approximately accommodate to a simple exponential decay (recall that the H2BGFP content dilutes two-fold with every cell division):

$$I(t) = I(0) \times 2^{-\lambda t} \tag{12}$$

where I(0) represents the initial cellular H2BGFP intensities. By contrast, if there were subpopulations of keratinocytes dividing at different rates (e.g. $\lambda_S << \lambda_P$), these would progressively segregate into different modes in the distribution of H2BGFP intensities at relatively long term, as $t \gg 1/(\lambda_P - \lambda_S)$. A first visual inspection at the experimental data revealed individual-keratinocyte H2BGFP intensities remained overall tightly distributed and scarcely dispersed even at latest 14d-18d time points across the different tissues (**Fig. 2B** and **Fig. S2B,D,F**). To formalize and automate the analysis, additional quantitative methods were adopted.

*Simulating H2BGFP dilution kinetics under different scenarios of cell proliferation*

An initial aim was to explore the quantitative H2BGFP dilution predictions and inference limits set by the different models of cell renewal under reasonable parameter assumptions (**Fig. 1D**). For this, we implemented stochastic simulations of the basal cell proliferation dynamics under each of these theoretical scenarios, in a similar manner as in **section 2**, with the peculiarity that simulations were embodied with real distributions of H2BGFP intensities I(0) in the initial cell populations and as individual cell division events occurred, these were linked with two-fold

H2BGFP partitioning (**Fig. 2D**). An additional noise term was included so that on average H2BGFP content in daughter cells differed by ~10%, in agreement with variability observed *in vitro*. Preliminary results using standard Markov-chain Monte-Carlo simulation methods (Gillespie's algorithm) indicated that for common average division rates $\lambda_P$ ~ 1.5–3/week, a two-to-three weeks chase would be enough to reliably distinguish a 10% subpopulation of basal stem cells dividing at a ≥4-fold slower rate $\lambda_S$ as a separate, retarded peak in the distribution of H2BGFP intensities.

A major issue of the Markov-chain Monte-Carlo implementations is that they assume kinetic processes are *memoryless*, and therefore, ignore the waiting times between consecutive cell divisions [11]. In other words, they consider that the probability that a cell divides in a time interval is independent of its current state and previous time spent in progressing through the cell cycle and only depends on the particular value of λ, so that, for a certain population, the time for completion of the cell cycle $t_{cc}$ satisfies an underlying exponential distribution:

$$P(t_{cc}) \sim \lambda e^{-\lambda t} \tag{13}$$

While this assumption may be acceptable to reproduce long-term dynamics, we found it had major limitations for a realistic description of cellular turnover at short-time scales, as for a given subpopulation with an average cell-cycle period $\langle t_{cc} \rangle = 1/\lambda$, at random, some cells would divide almost immediately after being born while some others would exhibit cell-cycle periods much longer than the average. For this reason, we extended the Monte Carlo algorithm to allow Non-Markovian simulations that permitted to explore alternative, more realistic cell-cycle time distributions (**Fig. 2D**; **Fig. S5A**). In particular, for each subpopulation cycling at a different average rate λ we implemented a delayed exponential distribution [12]:

$$P(t_{cc}) \sim \tau_R + Exp(\varphi) \tag{14}$$

where there would be a refractory period $\tau_R$ between consecutive cell divisions (i.e. a minimum cell-cycle time before keratinocytes can commit to divide again). In this case, $\langle t_{cc} \rangle = 1/\lambda = \tau_R + 1/\varphi$. However, more generally, we considered a whole family of hypothetical delayed Gamma distributions for the cell-cycle periods:

$$P(t_{cc}) \sim \tau_R + Gam(\kappa, \theta) \tag{15}$$

where κ and θ represent the shape and scale parameters of a Gamma distribution. Notice that for κ = 1, this distribution is a delayed exponential (i.e. Eq. 15 becomes equivalent to Eq. 14) and as κ gets larger, the cell-cycle period distribution would be assumed narrower. In this scenario, $\langle t_{cc} \rangle = 1/\lambda = \tau_R + \kappa\theta$.

The implementation of Gamma-shaped cell-cycle period distributions required modifying the time update process but also the initialization condition in the simulation code to allow for random, unbiased asynchronous cell cycle states at the starting time t0. The probability of capturing a cell at a particular time post-division τ would not be uniform but proportional to the probability of showing longer cell-cycle periods $t_{cc} > \tau$, and follows this equation:

$$P(\tau) = C \times \begin{cases} 1 & \tau < \tau_R \\ 1 - GamCDF(\tau - \tau_R, \kappa, \theta) & \tau \geq \tau_R \end{cases} \tag{16}$$

where C is a normalization factor and GamCDF is the particularized Gamma cumulative distribution function. By integrating Eq. 16, one can deduce that:

$$P(0 \leq \tau < \tau_R) = C \times \tau_R$$
$$P(\tau_R \leq \tau < \infty) = C \times \int_{\tau_R}^{\infty}(1 - GamCDF(\tau - \tau_R))d\tau \quad (17)$$

Accordingly, in the simulations, initial cells were assigned random cell-cycle states τ drawn from the corresponding underlying distributions (Eq. 16 and Eq. 17), so that a certain fraction were realistically ascribed to early stages of the cell-cycle ($\tau < \tau_R$) and thus required longer to undergo the first round of division (and therefore start the H2BGFP content dilution).

Following this refined methodology, we explored the H2BGFP dilution pattern predictions I(t) in different theoretical scenarios, including the SC-CP and 2xSC models under the corresponding parameter conditions set by [5], and [4]. Taking assumptions on the underlying cell-cycle time distributions in agreement with common estimates provided below (for quickly-dividing cells: $\tau_R$ = 0.5d, κ = 8; for slow-cycling cells: $\tau_R$ = 0.1x $\langle t_{cc} \rangle$, κ = 8), we found all these previous cases involving heterogeneous populations of slow- and quickly- dividing cells would lead to separated peaks in H2BGFP histograms by three weeks, in contrast to SP model predictions where individual-cell H2BGFP distributions would remain as a single peak (**Fig. S2A**).

*Unimodality tests and cell-proliferation model inference*

In order to formalize the classification of H2BGFP dilution patterns and test the efficiency to discriminate between homogeneous and heterogeneous proliferating cell population hypotheses, we next applied multiple statistical tests for unimodality. Six different methods were considered: Hartigan & Hartigan (1985) dip test (HH) [13], Silverman (1981) critical bandwidth test (SI)[14], Cheng & Hall (1998) excess mass test (CH) [15], Hall & York (2001) critical bandwidth test (HY) [16], Fisher & Marron (2001) Cramer-von Mises test (FM) [17] and Ameijeiras–Alonso et al (2018) excess mass test (ACR) [18]. All these tests yielded significant p-values for the H2BGFP distributions of the SC-CP and 2xSC scenarios set above, classifying them as multimodal (**Fig. S2A**).

Having demonstrated the reliability on synthetic data sets, we applied the unimodality tests to the experimental data to estimate the likelihood that the evolving fluorescence intensity distributions arose from a single proliferating cell population or cells dividing at multiple rates. Empirical distributions remained largely unimodal over time across body sites, both by analysing individual mice-derived data (**Fig. 2C**) or individual fields of view separately (**Table S3**). ~5 % of all samples were identified as multimodal based on p=0.05 (without multiple comparison correction). We found that there was no clear relationship between collection time and multimodality, and that, when observed, multimodality was inconsistent between animals. Furthermore, regardless of collection time, cell subpopulations classified as multimodal differed in a single division round, suggesting that the time of collection for those animals could occur during progression of synchronised divisions.

The analyses performed per field of view more strongly favoured unimodality than the individual animal measures, suggesting minor spatial asynchronies in the cell division timing (**Fig. S3C**). Altogether, our data are consistent with a single average rate of cell division λ at any given site, supporting the simplest SP model paradigm throughout the oesophagus and different skin niches.

*Estimating the distribution of keratinocyte cell-cycle times*

To further validate the SP model, we fit models of cell-cycle time distributions to the observed H2BGFP intensity profiles over time. If a unique mode of cell proliferation prevailed across the entire population of keratinocytes, the full shape of the cellular H2BGFP distributions at different times would be reproducible by means of a model distribution of cell-cycle periods (Eq. 15).

A grid-based Approximate Bayesian Computation (ABC) rejection method [19], was implemented to fit the time series of experimental H2BGFP intensity distributions with results from SP model simulations varying the values of the unknown cell-cycle parameters $\tau_R$ and $\kappa$. In this methodology for each tissue we fixed the value of the parameter $\lambda$ ($\lambda$ = 2.9, 2.0, 1.5, 1.2/week for oesophagus, hind-paw, ear and dorsum, respectively; estimated from the linear slope in the semi-logarithmic plot showing log(I) vs. time; see Eq. 12). Basal cell H2BGFP content dilution was simulated starting from initial values drawn from the corresponding empirical I(0) distribution, used as prior, and similar assumptions on label partitioning were considered as the ones described above.

A distance metric was computed for every $\{\tau_R, \kappa\}$ value pair based on the sum of absolute quantile differences between the simulated and the empirical I(t) histograms at the different time points, using quantiles taken at 0.025, 0.25, 0.5, 0.75, and 0.975. This identified a family of acceptable shapes for the cell-cycle period distribution, obtained as posterior estimates (**Fig. 2F**; **Fig. S2C,E,G**). In agreement with the SP paradigm, we obtained adequate fits on the whole series of keratinocyte H2BGFP dilution patterns with the following cell-cycle attributes: $\{\tau_R$ = 0.5, 1.0, 0.5, 0.5 days; $\kappa$ = 8, 4, 8, 16$\}$ for oesophagus, hind-paw, ear and dorsum, respectively (**Fig. 2B**; **Fig. S2B,D,F; Table S4**). Note using the outcome of a Kolmogorov-Smirnov test as an alternative distance metric did not substantially alter the cell-cycle solutions.

#### 4. Single-progenitor parameter inference from clonal data sets

Nine independent lineage-tracing data sets were exploited to challenge the suitability of the SP model to explain clonal dynamics in the esophagus and the different skin territories, and ultimately infer the most-likely parameter values of keratinocyte cell behavior. Given the diverse methodologies and disparity of inference results described so far in the literature (see **section 5**), we decided to undertake a single, robust, maximum likelihood estimation (MLE) approach for model fitting across data sets. In particular, a comprehensive grid search was performed on the unknown SP model parameters, and for every set of parameter values θ we run multiple simulations (see below) to get a theoretical estimate of the time course in the basal clone size distributions, to be contrasted with the experimental one. A log-likelihood value $l(\theta; x)$ was calculated as follows:

$$l(\theta; x) = \sum_t \sum_n (x_n(t) * \log p_n(t, \theta)) \quad (18)$$

where $x_n(t)$ is the observed frequency of clones of a certain basal size *n* at time *t*, and $p_n(t, \theta)$ is the probability of observing clones of that size at time $t$ given the parameter values θ, a quantity obtained from the model simulations.

Note that, given the particular scaling behaviour of the clone size distributions and their large asymmetry (approximately log-normal-like shapes), clone sizes were conveniently binned in ranges increasing in powers of two, as done in previous work [10], i.e. $n$ above stands for clones with a number of basal cells in the range ($2^{n-1}$ + 1) to $2^n$. Also, only surviving clones with at least 2 basal cells were considered for the MLE analysis, to exclude any possible contribution due to post-mitotic cells labelled at induction (recall the initialization condition for simulations was $P_n(t0) = \delta_{n_P,1}\delta_{n_D,0}$). Additionally, in some experimental data sets a small proportion (<1%) of late-time clones were reported with sizes that greatly exceeded the vast majority of their coexisting clones (i.e. sizes >> 2.3*SD above the mean clone size, a reference for the 99% CI threshold of log-normal distributions). This occurred for 3, 4 and 5 clones in the data sets from esophagus [20], dorsum [10] and paw [1], respectively [20]. These outlier clones, which could be interpreted as a result of non-neutrality or coincidental fusion between two adjacent clones, were pooled together and assigned into the immediate prior category of clone sizes to circumvent the computational issues of estimating the probability of extremely rare events with sufficient precision. However, excluding these outstandingly large clones from the analysis did not substantially alter the parameter estimates provided below.

Maximum likelihood estimates $\hat{\theta}_{MLE}$ (maximizing the expression in Eq. 18) were obtained for each data set, and presented with 95% confidence intervals computed based on the likelihood-ratio test [21]. Parameter solutions were plotted as heatmaps in 2D or 3D parameter spaces (as applicable), color coded according to the value of the log-likelihood ratio statistic (the maximum value of 0 corresponding with the $\hat{\theta}_{MLE}$). Parameter sets with values falling below -7.81 (the $\chi^2$ statistic cutoff for an α=0.05, 3 degrees of freedom) were considered non-optimal and generally not displayed.

*Cell-cycle time -constrained parameter inference*

As a first approximation to parameter inference, simulations were performed using Gillespie's Markov-chain Monte-Carlo algorithm (as described in **section 2**) and all the 3 parameters of the SP model were considered unknown (**Fig. S4A**). We thus iterated on values of $r \in [0, 0.5]$, $\rho \in [0, 1]$ and $\lambda$ (within a reasonable range of values: $\in [0.4, 3.6]$/week) (recall ρ = Γ / (Γ + λ) in homeostasis), testing a total of ~230,000 parameter conditions in a 3D space (super-computing resources at the Wellcome Sanger Institute were used for parallelization). Multiple parameter combinations yielded relatively good fits on clone size distributions over time (**Fig. S4B**). λ may be accurately measured independently by H2BGFP dilution experiments (**section 3**), so that, in practice, we used this estimate to constrain our parameter search to 2D (a total of 101 × 100 parameter combinations were explored). By this means, we increased the discrimination on the values of the parameters *r* and ρ (or Γ), yet a certain level of imprecision remained among solutions aligning around constant *r*/ρ ratios, leaving a characteristic, relatively long trail of coloured patterns in the corresponding heatmaps (**Fig. S4**) (see below).

One could potentially speculate if this level of parameter imprecision was due to biological variability (e.g. inter-mice differences or age-related differences in cell behaviour across the distinct time points). However, we separated the MLE calculation into individual time point analyses, observing each was consistent with a similar pattern of degenerated solutions (**Fig. S4C**). We also generated synthetic data sets by strict simulation of the SP model under specific parameter values and submitted their clone size distributions to a similar MLE inference analysis, obtaining comparable levels of inaccuracy for same sample sizes (**Fig. S4C**). This suggested that parameter uncertainty was not biological, nor due to a flawed, inappropriate SP-model definition. We highlight in the different experimental heatmap panels how the direction in our MLE parameter uncertainties fell indeed consistent with the *r*/ρ ratios obtained from the asymptotic linear slope in the average clone size over time (**Fig. 3D**, grey lines) (dashed lines correspond with the 95% CI limits in the linear slope). It follows that it is an inherent feature of the stochastic nature of clone fates and their quick convergence into a scaling form, as extended synthetic data sets revealed: the level of imprecision could in theory be further attenuated by larger experimental sample sizes at relatively early time points (**Fig. S4D**).

A second issue arises from the possible impact the assumptions on the cell-cycle time distribution may have on clone-size estimates and hence on parameter inference. For that reason, Non-Markovian simulations of the SP model were tested, with different hypothetical underlying cell-cycle time distributions, as we did in **section 3** for cell-proliferation studies (**Fig. S5A**). Theoretical simulation results confirmed clone size frequencies predicted at relatively early time points differed substantially between implementations carried out with Gamma-distributed cell-cycle periods and those with default exponential assumptions, differences getting smaller over time, as shown by Kullback–Leibler divergence (**Fig. S5B-C**). On average, it was not until a critical time $T_c$ of 3 (2; 10) × $\langle t_{cc} \rangle$ that details of the shape of cell-cycle time distribution became unimportant on clonal predictions. This meant the shortest experimental time points, when clonal data have not yet fully converged to the long-term scaling behavior and can potentially improve the precision of estimated cell parameters, were also more prone to contribute to a biased inference given unrealistic assumptions on the cell-cycle time distribution (**Fig. S5D-E**). Therefore, our SP model simulations used for MLE parameter inference were constrained for each body site by the actual cell-cycle time distribution estimated from the corresponding H2BGFP dilution analysis (**section 3**).

Following this more realistic methodology, the SP parameter estimates $\hat{\theta}_{MLE}$ generally shifted towards lower values of ρ (and a slower stratification rate Γ) than those predicted with default Markovian simulations (i.e. exponential $t_{cc}$ distributions), discarding hypothetical scenarios where differentiating cells would be largely absent in the basal compartment and would stratify almost immediately after being born (**Fig. 3D**). Excellent fits were obtained on the time courses of the experimental clone size distributions --including early and long-term clonal behavior-- across data sets, resulting in a statistically significant improvement over the model predictions made by the original publications (**Fig. 3E**; **Fig. S8A-C; Table S4**). Compared to our $\hat{\theta}_{MLE}$, the value of the log-likelihood ratio statistic of previous estimates was: -9.4 (for Doupé et al's [20] fits in esophagus), -364.2 (for Lim et al's [1] fits in hind-paw epidermis), -29.3 (for Doupé et al's [22] fits in ear epidermis) and -13.0 (for Murai et al's [10] fits in dorsal epidermis). Altogether, our fits confirm the suitability of the SP model and provide more accurate descriptions of the parameters defining keratinocyte cell behavior in the different territories (**Table 1**).

#### 5. Revisiting alternative datasets and interpretations

Here we revisit published work from the literature in order to test the ability of a cell-cycle time-sensitive SP model to explain these different datasets, and we reexamine the specific arguments and claims previously given in support of alternative models.

##### *Giroux et al (2017), esophageal epithelium*

Giroux et al [23] performed lineage tracing in mouse esophageal epithelium using a *Krt15* promoter, and postulated the existence of a long-lived subpopulation of stem cells, characterized by high expression of Krt15. The authors concluded on the heterogeneous proliferation potential of the esophageal basal cells –a scenario compatible with a hierarchical stem-cell transit-amplifying cell model – based on the molecular properties of the Krt15$^{+}$ basal cells and the long-term persistence of a subset of *Krt15*-labelled clones well beyond the homeostatic renewal time of the epithelium, giving rise to all differentiated lineages. This observation is however consistent with the SP model, since, as noted above, a small number of clones dominate the tissue after extended periods. Giroux et al [23] did not report clone sizes, but data on clonal densities could be recovered from presented figures. Using these experimental lineage-tracing results in *Krt15-Cre$^{PR1}$ R26$^{mT/mG}$* mice as an independent dataset, we found that the single progenitor model proposed for the esophagus was indeed capable of reproducing observed data. The best parameter estimates obtained from the analysis of *Lrig1-eGFPcre$^{ERT}$ R26$^{flConfetti}$* clones produced excellent fits on the time course in the *Krt15*-labelled clone density from [23] (**Fig. S7D; left panel; Table S4**). Finally, we extended our analysis to consider the label distribution across differentiated cell layers (Fig. 2E in [23]. Again, we observed the SP predictions were consistent with experimental observations (**Fig. S7D; right panel; Table S4**), indicating that a hierarchy is not required to explain esophageal epithelium dynamics.

##### *Mascré et al (2012) and the SC-CP model*

Mascré et al [5]propose a proliferative hierarchy of slow-cycling stem cells underpinning committed progenitor cells (SC-CP model) from the quantitative analysis of clonal fates in the tail interfollicular epidermis. The authors argue that cells labelled with two different inducible genetic constructs targeting the promoters *Ivl* and *Krt14* have distinct dynamics. They conclude that *Ivl* and *Krt14* are markers of P cells and both P and S cell populations, respectively. However, in this paper the authors did not test the ability of alternative models to describe the observed data. The SC-CP model was explicitly claimed to explain the diverging trends in the surviving clone fraction and the labelled cell fates at the earliest time points.

Unfortunately, the clonal data was not available. Alternatively, cell proliferation-related data could be extracted from plots of an independent H2BGFP dilution experiment performed to validate their predictions. We reanalyzed the displayed distributions of the number of cell divisions (see Fig. 3k in [5]) and found that a SP model ($\lambda$ = 1.3/week; cell-cycle distribution with $\tau_R$ = 0.6 days, $\kappa$ = 1.5) (**Table S4**) could provide a similar, suitable fit on the experimental data along the different chase times (**Fig. S10A**). Furthermore, no bimodality was observed in the distribution of histone intensities, as one might expect by the 3 week timepoint if there was a significant subpopulation of slow-cycling stem cells dividing at $\lambda_S \approx 0.1$/week (**Fig. S2A; section 3**).

A limitation for this study is that the structural heterogeneity of murine tail was not considered, when distinct spatial territories (scale and interscale regions; **Fig. S3A**) are believed to show different developmental processes, cell proliferation rates, and differentiation programs[24]. As in [3], this spatial information was not considered in [5], where H2BGFP fluorescence was analyzed from FACS on pools of basal ($\alpha_6$-integrin$^+$ CD34$^-$) cells. Another potential issue was that no labelling and exclusion of immune, CD45$^+$ cells was performed in the study of label retaining cells, which could introduce a source of error.

*Sánchez-Danés et al (2016), tail skin*

As a continuation to the work in Mascré et al [5] [6] revisited clonal dynamics in tail skin by inducible genetic labeling using same targeted promoters, *Ivl* and *Krt14*, but analyzing labelled clones independently in scale and interscale regions. While they conclude that a SP model explains clonal dynamics in scale, they argue the SC-CP model prevails in interscale on the basis of the different clonal dynamics observed using the *Ivl-Cre*$^{ER}$ and *Krt14-Cre*$^{ER}$ constructs. The evidence used to make this argument is an apparent decrease in the labelled cell fraction over time for the *Ivl-Cre*$^{ER}$-targeted population (argued to only label committed progenitors), and an increase in the labelled cell fraction of the *Krt14-Cre*$^{ER}$-targeted clones (considered to comprise driving stem cells) (see Fig. 2e in [6]) (Eq. 10). Here, the labelled cell fraction was *estimated* as a product of the average basal clone size and the overall clone density, which is discussed later.

For the purpose of model fitting and validation they assumed division rates for the P cells similar to those reported in [5] in both scale or interscale regions ($\lambda_P \approx 1.2$/week). A least-squares minimization procedure was then used to fit the evolution of the mean clone sizes for each construct in each compartment, using the corresponding ascribed model, either the SP or the SC-CP model. They found the best fit for the labelled *Krt14-Cre*$^{ER}$ clones in interscale was attained with a SC-CP model where $\lambda_S = 0.45$/week, $r_S = 0.03$, $\lambda_P = 1.7$/week, $r_P = 0.19$, $\Delta = 0.02$ and $\chi^{div}_{S,label} = 0.65$ (see Fig. 2d in [6]). The confidence intervals in the value of $\Delta$ span the $\Delta = 0$ condition, suggesting that a fit on clonal dynamics would be possible without P cells showing a necessary imbalance towards terminal differentiation, raising the question of whether a SP model could recapitulate the data.

We therefore fitted the experimental basal clone size distributions [6]with a SP model, taking $\lambda_P = 1.2$/week, as in [5]. The best-fit parameter sets showed improved fits on both the scale- and interscale- *Krt14-Cre*$^{ER}$ clone data (**Fig. S10B; Table S4**). Our results for interscale demonstrate that a SP model ($\lambda_P = 1.2$/week, $r = 0.09$, $\Gamma = 2.2$/week) showed satisfactory fittings on the clone size frequencies, average clone size and clonal survival over time (**Fig. S10B-D**).

Given that time courses in clone size and clonal survival are both consistent with the SP model, it is sensible to consider whether the apparent trends in the labelled cell fraction can also be explained within this paradigm. While the labelling of a large, representative set of dividing cells in homeostasis would in principle remain overall constant over time, there are two possible sources of variation that could contribute to the observed deviations with the SP model: stochastic growth or decline in a finite labelled population, and inter-animal variation in initial label induction. To address the question of whether these sources of random variation could be sufficient to explain the extent of increase in the *Krt14-Cre*$^{ER}$- labelled cell fraction, we reexamined noise by *error propagation* in both the original data and simulated SP model.

To account for inter-animal variation in label efficiency, we measured SD from clonal density data by exclusively studying a single, arbitrarily selected sub-region per animal ($N$ = 2-5 mice per time point). In the original study, multiple distinct sub-regions were treated as independent observations, reducing the apparent error arising from variable labelling. The SD in clone density was then integrated together with the SEM in basal clone size to obtain the experimental error in the *estimated* label cell fraction at each time. To estimate variations in average clone size due to SP stochasticity, we subsequently run time-course simulations tracking the same number of clones counted in the experiments ($N$ = 72, 75, 40, 31, 47, 70 clones sampled at time 1, 2, 4, 8, 12 and 24w, respectively), and measured the SEM and average clone size obtained through multiple runs. This info was combined with independent time-course simulations of clone density reproducing the variable clonal induction. From this analysis we find that any trend in the experimental *Krt14-Cre$^{ER}$*-labelled cell population from [6] largely fell within the domain of uncertainty given by the combined sources of variation, considering the actual level of sampling error, with just the very last time point being at the borderline of the 95% confidence interval **(Fig. S10E)**.

Finally, given that *Krt14-Cre$^{ER}$* clonal dynamics can be explained by the SP model, it raises the question of how distinct the *Ivl-Cre$^{ER}$* interscale clone population behavior is. Indeed, excluding the earliest two experimental time points (up to 2 weeks post-induction, where dynamics could be potentially influenced by initial priming of labelled cells towards differentiation; recall Eq. 11), we find an adequate fit over the time courses in the average clone size, clonal survival and labelled cell fraction with just the same parameter values obtained from fitting *Krt14-Cre$^{ER}$* populations (**Fig. S10F-H; Table S4**). This suggests that two distinct promoters could target the same, unique proliferative cell type.

*Sada et al (2016) and the 2xSC model*

Sada et al [4] propose a model of epithelial cell renewal where tissue is maintained by two independent populations of dividing stem cells cycling at different rates (2xSC model). This model was based upon observations made in H2B-GFP dilution experiments in back-skin keratinocytes. In these experiments the authors tracked H2B-GFP histograms of FACS-sorted epidermal cells from *Krt14-cre$^{ERT}$ R26$^{tdTomato}$ Krt5$^{tTA}$/pTRE-H2BGFP* mice collected after different chase times with doxycycline (**Fig. S9A**).

Based on this experiment, the authors reported resolved distributions of the number of cell divisions for the basal, spinous and granular layers separately. They further attempted to fit different published models to the data, finding that neither the SP-model nor the SC-CP model adequately described the data, whilst the 2xSC model was compatible (see Fig. 3 in [4]). Here we examine their model *implementation* (in contrast to the more general mathematical description given in **section 2**).

In their model, the authors considered two subpopulations of basal cells, $S_1$ and $S_2$. Cells in the $S_2$ population may either divide or stratify. Division occurs slowly, giving two $S_2$ daughter cells, and stratification is conceived as an independent, "division-uncoupled" process. That is to say, cell fate is not determined on birth. We note that this subpopulation would in fact behave equivalently to a constrained SP-model where r=0.25 (e.g. see [12])[2]. Regardless of whether

[2] Authors' decision to consider $S_2$ basal cells as a single pool undergoing symmetric divisions is perhaps arbitrary, since in homeostasis half of these cells should go on to divide and half should stratify (i.e. can be called D-cells), hence the reason for r=0.25 using the SP-model formulation (Eq. 1).

biological differentiation initiates prior to, concomitantly to, or after stratification, a class of basal "D" cells (ignored by the authors) arising from the $S_2$ population can be defined *post hoc,* as those cells that proceed to stratify (as described above in **section 2**). Thus, the time of fate determination does not alter the model formulation and $S_2$ cells can be considered as a stochastic SP population (Eq. 1).

$S_1$ cells in contrast divide quickly and asymmetrically, giving one $S_1$ daughter and one stratifying, suprabasal daughter, implying any differentiating D cell stratifies immediately upon birth ("division-coupled stratification"). This is a deterministic process of invariant asymmetric self-renewal, and as such, would not be supported by clone size distributions observed in lineage tracing experiments (see **section 2**).

To resolve this issue, the authors introduce a variant of the 2xSC model (termed the "hybrid" 2xSC model, as opposed to the former "semi-coupled" one) where the $S_1$ population undergoes a combination of symmetric and asymmetric divisions. It follows that $S_1$ differentiating cells should stochastically choose between waiting and following an uncoupled stratification process (with probability $u_{S1}$) or stratifying immediately after division (with probability 1- $u_{S1}$). We note that whilst this new complex mechanism could reproduce clone size distributions, there exists no direct evidence to support it.

The hybrid 2xSC model was directly implemented using the following differential equations to account for H2B-GFP dilution kinetics (pg.14-16 in Supplementary Note in [4])[3]:

$$\begin{aligned} D_t\eta_d^{S1}(t) &= (1+u_{S1})\lambda_{S1}\eta_{d-1}^{S1}(t) - (\lambda_{S1} + u_{S1}k_{S1\to SL})\eta_d^{S1}(t) \\ D_t\eta_d^{S2}(t) &= 2\lambda_{S2}\eta_{d-1}^{S2}(t) - (\lambda_{S2} + k_{S2\to SL})\eta_d^{S2}(t) \\ D_t\eta_d^{SL}(t) &= u_{S1}k_{S1\to SL}\eta_d^{S1}(t) + (1-u_{S1})\lambda_{S1}\eta_{d-1}^{S1}(t) + k_{S2\to SL}\eta_d^{S2}(t) - k_{SL\to GL}\eta_d^{SL}(t) \end{aligned} \quad (19)$$

subjected to boundary conditions $\eta_d^x(0) = \delta_{d0}\rho_x$, where $\eta_d^x(t)$ represents the density of cells of type *x* that have divided *d* times by time *t.* In the extreme, semi-coupled scenario ($u_{S1}$ = 0), the equations were reported as:

$$\begin{aligned} D_t\eta_d^{S1}(t) &= \lambda_{S1}\left(\eta_{d-1}^{S1}(t) - \eta_d^{S1}(t)\right) \\ D_t\eta_d^{S2}(t) &= 2\lambda_{S2}\eta_{d-1}^{S2}(t) - (\lambda_{S2} + k_{S2\to SL})\eta_d^{S2}(t) \\ D_t\eta_d^{SL}(t) &= \lambda_{S1}\eta_{d-1}^{S1}(t) + k_{S2\to SL}\eta_d^{S2}(t) - k_{SL\to GL}\eta_d^{SL}(t) \end{aligned} \quad (20)$$

Here, we reimplemented Sada et al's model to test its properties. To do so, we first displayed the hybrid 2xSC model mathematically, in terms of its constituent processes and associated transition probabilities:

$$S_1 \to \begin{cases} \xrightarrow{\lambda_{S1}} S_1 + S_1 & \text{Prob. } u_{S1}/(1+u_{S1}) \\ \xrightarrow{\lambda_{S1}} S_1 + SL & \text{Prob. } (1-u_{S1})/(1+u_{S1}) \\ \xrightarrow{k_{S1\to SL}} SL & \text{Prob. } u_{S1}/(1+u_{S1}) \end{cases}$$

[3] In a later refinement, the authors replaced the $S_1$ and SL processes by two-step Poisson processes involving two subpopulations, each having twice the rate of the original process, in order to recreate Gamma waiting-time distributions (κ=2). This final model was the one used to fit to the experimental data, and thus the one we reassessed. Nevertheless, our main points below arise from the core implementation, and apply irrespective of waiting-time considerations, reason why we present equations as in their simple version for clarity.

$$S_2 \rightarrow \begin{cases} \xrightarrow{\lambda_{S2}} S_2 + S_2 & \text{Prob. } 0.5 \\ \xrightarrow{k_{S2\rightarrow SL}} SL & \text{Prob. } 0.5 \end{cases} \quad (21)$$
$$SL \xrightarrow{k_{SL\rightarrow GL}} \emptyset$$

where $\lambda_{S1}$, $k_{S1\text{->}SL}$, and $\lambda_{S2}$, $k_{S2\text{->}SL}$ denote the division and stratification rates of the $S_1$ and $S_2$ populations, respectively. SL refers to the population of cells in the first suprabasal (spinous) compartment, which would transit with rate $k_{SL\text{->}GL}$ to the granular layer and to more external layers successively (here not depicted). In homeostasis, $k_{S1\text{->}SL} = \lambda_{S1}$, $k_{S2\text{->}SL} = \lambda_{S2}$, and $k_{SL\rightarrow GL} = \left(\lambda_{S1}\chi_{S1}^{div}/(1+u_{S1}) + 0.5\,\lambda_{S2}\left(1-\chi_{S1}^{div}\right)\right)/m$, where $\chi_{S1}^{div}$ is the fraction of $S_1$ basal cells and $m$ is the spinous suprabasal-to-basal cell ratio. The event probability terms in Eq. 21 are those required for homeostasis. From Eq. 21 it becomes apparent how Sada et al's hybrid 2xSC model represents a particular case of a two SP model combination where $r_{S2} = 0.25$, $r_{S1} = u_{S1}/(1+u_{S1})$ and $\Gamma \rightarrow \infty$ (i.e. differentiating D-cells stratifying immediately upon birth to yield SL cells). That is why in **section 2** we adopted the more general form to represent the 2xSC model (Eq. 2) (in that case we referred to the slow-cycling population as $S_1$ for convenient comparison with SP and SC-CP schemes).

When we derive the kinetic equations for H2B-GFP dilution analysis from Eq. 21 we obtain:

$$\begin{aligned} D_t\eta_d^{S1}(t) &= \lambda_{S1}\eta_{d-1}^{S1}(t) - \left(\tfrac{1}{1+u_{S1}}\lambda_{S1} + \tfrac{u_{S1}}{1+u_{S1}}k_{S1\rightarrow SL}\right)\eta_d^{S1}(t) \\ D_t\eta_d^{S2}(t) &= \lambda_{S2}\eta_{d-1}^{S2}(t) - 0.5(\lambda_{S2} + k_{S2\rightarrow SL})\eta_d^{S2}(t) \\ D_t\eta_d^{SL}(t) &= \tfrac{u_{S1}}{1+u_{S1}}k_{S1\rightarrow SL}\eta_d^{S1}(t) + \tfrac{1-u_{S1}}{1+u_{S1}}\lambda_{S1}\eta_{d-1}^{S1}(t) + 0.5k_{S2\rightarrow SL}\eta_d^{S2}(t) - k_{SL\rightarrow GL}\eta_d^{SL}(t) \end{aligned} \quad (22)$$

For $u_{S1} = 0$ (semi-coupled case), these simplify to:

$$\begin{aligned} D_t\eta_d^{S1}(t) &= \lambda_{S1}\left(\eta_{d-1}^{S1}(t) - \eta_d^{S1}(t)\right) \\ D_t\eta_d^{S2}(t) &= \lambda_{S2}\eta_{d-1}^{S2}(t) - 0.5(\lambda_{S2} + k_{S2\rightarrow SL})\eta_d^{S2}(t) \\ D_t\eta_d^{SL}(t) &= \lambda_{S1}\eta_{d-1}^{S1}(t) + 0.5k_{S2\rightarrow SL}\eta_d^{S2}(t) - k_{SL\rightarrow GL}\eta_d^{SL}(t) \end{aligned} \quad (23)$$

Comparing Eq. 22-23 with Eq. 19-20, we observe that Sada et al omitted some probability terms in their equations; most noticeably the 0.5 factor accompanying the $\lambda_{S2}$ parameter, which could have potentially impacted on the estimates given for $S_2$ the division rate. To shed light on this, we simulated H2BGFP dilution kinetics of the 2xSC model, taking the estimated parameter values from in the paper [4]: $\lambda_{S1}$ = 0.47/day, $\lambda_{S2}$ = 0.19/day, $u_{S1}$ = 0.20, $\chi_{S1}^{div}$ = 0.74 [4]. We found that we were unable to reproduce the reported fittings on the distributions of cell division number. In fact, these conditions resulted in bimodal H2BGFP dilution profiles and very poor fits against their experimental data (**Fig. S9B**: dashed blue lines) (This bimodality becomes even more prominent when considering narrower cell-cycle time distributions comparable to those we find in our experiments (**Fig. S2A**)). In contrast, when keeping all other constants equal but using $\lambda_{S2}$ = 0.38/day (a value twice the one reported), we found that the model fitted the data well, consistent with the fits originally presented by the authors, suggesting that the reported value for $\lambda_{S2}$ was incorrectly half of the actual rate (**Fig. S9B**: solid blue lines). To corroborate these findings, our analysis was performed both by stochastic simulation of processes in Eq. 21 (following the methods described in **section 3**) and by numerical integration of Eq. 22-23, obtaining the same results.

In light of the apparently similar rates of division for the $S_1$ and $S_2$ populations ($\lambda_{S1} \approx 3.3$/week and $\lambda_{S2} \approx 2.7$/week), we reinvestigated whether the SP model could provide an adequate fit of the data. Whilst the authors reported poor fits with a SP model (**Fig. S9B**: dashed orange lines), this model was just tested assuming rates could be adequately modelled as Poisson processes, whereas the 2xSC model was implemented using Gamma distribution waiting-time processes. As explored in **section 3**, the choice of distribution for cell-cycle times can substantially alter fits for histone dilution experiments, and we therefore took account of this explicitly.

Using the methods described in **section 3**, we explored different rates and waiting time values for the SP model and obtained fits similarly good as with the 2xSC model with a single population with division rate $\lambda$ = 3.1/week with a minimum refractory period $\tau_R$ = 0.4 days, and a stratification rate $\Gamma = k_{SL\text{->}GL}$ = 16x $\lambda$ following a Gamma distribution with shape parameter $\kappa_{strat}$ = 2.8 (**Fig. S9B**: solid orange lines; **Table S4**). We therefore conclude that the SP model proves suitable to explain the dataset in [4].

*Füllgrabe et al (2015), back skin*

Füllgrabe et al [25] performed genetic lineage tracing followed by quantitative analysis of clone dynamics from *Lgr6*-expressing back-skin cells, observing scaling properties indicative of stochastic, population asymmetry self-renewal in the interfollicular epidermis of murine dorsum [25]. Although the authors did not attempt a detailed model verification/parameterization, we considered the lineage tracing outcome from *Lgr6-cre$^{ERT}$/R26$^{Confetti}$* mice could constitute an ideal independent dataset to challenge our SP-model predictions against. Given the fact that labelling of $Lgr6^+$ cells was induced early in postnatal development (P3w), at a time when the tissue was likely not yet homeostatic (see Fig. 1D in [25]), we restricted the analysis to the late time points where mice were at least 8w old and the tissue was representative of adulthood. To do so, we randomly sampled the observed clone sizes at 40 days post-induction as initial modelling condition and simulated clonal dynamics over the period up to the 100-days and 150-days timepoints using a SP model with the parameters inferred from the H2B-GFP pulse-chase (**Fig. S2F-G**) and *Ah-cre$^{ERT}$/R26$^{EYFP}$* system (**Fig. S8C**; **Table 1; Table S4**). To take account of the unknown composition of initial preformed clones, we randomized (according to a binomial distribution) the proportion of P-cells in each simulated initial clone according with the parameter $\rho$ = 0.61.

Our fits fell in agreement with the observed growth in the average clone size as well as with the experimental time courses in the frequencies of clone sizes (**Fig. S9C**). These results reaffirm the validity of the SP paradigm in murine back skin with the parameter values we calculated previously, using an independent genetic construct.
